# Lipid nanodisc scaffold and size alters the structure of a pentameric ligand-gated ion channel

**DOI:** 10.1101/2022.11.20.517256

**Authors:** Vikram Dalal, Mark J. Arcario, John T. Petroff, Noah M. Dietzen, Michael J. Rau, James A. J. Fitzpatrick, Grace Brannigan, Wayland W. L. Cheng

**Author notes:** authors contributed equally to this manuscript (Vikram Dalal, Mark J. Arcario, John T. Petroff). To whom correspondence should be addressed: Professor Wayland W. L. Cheng, Department of Anesthesiology, Washington University School of Medicine, MSC 8054-0043-12, Saint Louis, MO 63110.

## Abstract

Lipid nanodiscs have become the standard reconstitution system for structural and biochemical studies of membrane proteins, especially using single particle cryo-EM. We find that reconstitution of the pentameric ligand-gated ion channel (pLGIC), *Erwinia* ligand-gated ion channel (ELIC), in different nanodisc scaffolds (MSP1E3D1, SMA, saposin, spMSP1D1) produces distinct apo and agonist-bound structures. In the presence of agonist, different nanodiscs scaffolds produce concerted conformational changes associated with activation in ELIC, with larger nanodiscs showing more activated conformations. The effect of different nanodisc scaffolds on ELIC structure extends to the extracellular domain and agonist binding site. Molecular dynamic simulations of ELIC in small and large nanodiscs suggest that the impact of the nanodisc on ELIC structure is influenced by nanodisc size. Overall, the results indicate that the nanodisc profoundly affects the structure of a pLGIC, and suggest that larger circularized nanodiscs may be advantageous to approximate a lipid membrane environment.

## Introduction

Lipid nanodiscs are routinely used for the reconstitution of membrane proteins for structure determination by single particle cryo-EM. Multiple nanodisc scaffolds are available including membrane scaffold proteins (MSPs)^1^, saposin^2^, the synthetic co-polymer styrene maleic acid (SMA)^3^, and circularized MSP scaffolds^4,5^. While it is often assumed that nanodiscs mimic the environment of a model lipid bilayer, studies of the bilayer properties of empty MSP nanodiscs (i.e. nanodiscs without a membrane protein) indicate that nanodisc size has complex effects on membrane thickness, order and stiffness^6,7,8,9^. Furthermore, a recent cryo-EM structure of SARS-CoV-2 ORF3a in nanodiscs showed direct interactions of the scaffold with the membrane protein^10^. Therefore, it is possible that the nanodisc scaffold can influence membrane protein structure by altering the properties of the lipid bilayer or directly interacting with the protein of interest. Understanding this is essential to define functionally-relevant conformations and to characterize membrane effects on protein structure.

Structures of pentameric ligand-gated ion channels (pLGICs) in lipid nanodiscs have revolutionized the structural biology and pharmacology of these ion channels^11^. However, pLGICs are sensitive to their membrane environment^12,13^, which suggests that interactions between the nanodisc and protein might affect the conformation. Structures of several pLGICs have been determined in different nanodisc scaffolds including the glycine receptor (GlyR) in MSP1E3D1 and SMA nanodiscs^14^, and the GABA(A) receptor (GABA_A_R)^15,16^ and muscle-type nicotinic acetylcholine receptor (nAchR)^17,18^ in MSP2N2 and saposin nanodiscs. In the cases of GlyR and GABA_A_R, distinct conformations were observed in the different scaffolds. However, in both cases, different lipids or protein constructs were used, such that the impact of the scaffold per se on protein structure was unclear. Here, we report differences in the structure of the model pLGIC, ELIC (*Erwinia* ligand-gated ion channel), using different nanodisc scaffolds, and perform MD simulations to explore the underlying mechanism. The results indicate that nanodiscs scaffold and size alter the structure of pLGICs, and suggest that circularized nanodiscs may be the best scaffold to approximate a native-like membrane environment.

## Results

### Nanodisc scaffolds alter the structure of ELIC

We previously showed that ELIC in liposomes composed of 2:1:1 POPC (1-palmitoyl-2-oleoyl-phosphatidylcholine): POPE (1-palmitoyl-2-oleoyl-phosphatidylethanolamine): POPG (1-palmitoyl-2-oleoyl-phosphatidylglycerol) phospholipids (hereafter called 2:1:1 lipids) shows greater agonist responses than in POPC-only liposomes^19^. The peak open probability of ELIC in response to saturating concentrations of agonist in HEK membranes has been estimated to be >0.96^20^. Thus, it is plausible that the gating efficacy of ELIC in 2:1:1 lipids, which mimics the abundance of zwitterionic and anionic phospholipids in a cell membrane, is also quite high. One might therefore expect that an agonist-bound structure of ELIC in nanodiscs using these lipids should show a desensitized conformation. However, cryo-EM structures of agonist-bound ELIC in MSP1E3D1 nanodiscs using 2:1:1 lipids or only POPC show identical structures most consistent with a pre-active conformation^19,21^. We hypothesized that the nanodisc is altering the conformational landscape of ELIC, producing a discrepancy between the functional measurements in liposomes and structures in nanodiscs.

To test the impact of the nanodisc scaffold on ELIC structure, we reconstituted ELIC in three different nanodisc systems: styrene maleic acid (SMA)^3^, saposin^2^, and the circularized nanodisc scaffold, spMSP1D1^5^, using 2:1:1 lipids. The “sp” in spMSP1D1 refers to the Spycatcher-Spytag technology used to generate the circularized scaffold. Single-particle cryo-EM of ELIC in these nanodiscs in the presence of 50 mM propylamine yielded structures with an overall resolution of 3.96 Å for SMA, 3.50 Å for saposin, and 3.32 Å for spMSP1D1 (Supplementary Table 1, Supplementary Fig. 1 and 2). In a previous study, we reported structures of WT ELIC in MSP1E3D1 nanodiscs (i.e. a conventional linear scaffold) with 2:1:1 lipid in the absence and presence of agonist which are consistent with putative resting and pre-active conformations, respectively, as well as a structure of a non-desensitizing mutant, ELIC5 (P254G/C300S/V261Y/G319F/I320F), which is consistent with an open-channel conformation^19^. These structures will be referred to as apo-MSP1E3D1_ELIC_ (WT ELIC without agonist in MSP1E3D1), MSP1E3D1_ELIC_ (WT ELIC with agonist in MSP1E3D1) and ELIC5 (ELIC5 with agonist in MSP1E3D1) (Fig. 1). In this study, we compared the agonist-bound structures of ELIC in SMA, saposin, and spMSP1D1 (hereafter called SMA_ELIC_, saposin_ELIC_ and spMSP1D1_ELIC_) with these putative resting (apo-MSP1E3D1_ELIC_), pre-active (MSP1E3D1_ELIC_), and open-channel conformations (ELIC5) (Fig. 1). The diameter of the cryo-EM density for the nanodisc in each structure was also estimated from low-pass filtered maps as previously described^15^, producing diameters of 9.3 nm for MSP1E3D1_ELIC_, 10.2 nm for saposin_ELIC_ and 10.8 nm for spMSP1D1_ELIC_ (Fig. 1). The nanodisc structure for SMA_ELIC_ was too disordered to measure. Thus, among agonist-bound WT structures, varying the nanodisc scaffold altered the apparent size of the nanodisc density. MSP1E3D1_ELIC_ produced the smallest nanodisc density (9.3 nm), similar to the GABA_A_R in MSP2N2 nanodiscs^15^, and spMSP1D1_ELIC_ produced the largest nanodisc diameter consistent with the predicted size of empty spMSP1D1 nanodiscs (~11 nm)^5^. Interestingly, the diameter of the nanodisc density of apo-MSP1E3D1_ELIC_ is larger than MSP1E3D1_ELIC_, and ELIC5 (in MSP1E3D1 nanodiscs) is similar to spMSP1D1_ELIC_. We also assessed nanodisc size using size exclusion chromatography (Supplementary Fig. 3). The effective diameter of ELIC in MSP1E3D1, saposin and spMSP1D1 nanodiscs, which depends partially on nanodisc size, agrees with the trend observed in the structures.

**Figure 1:**
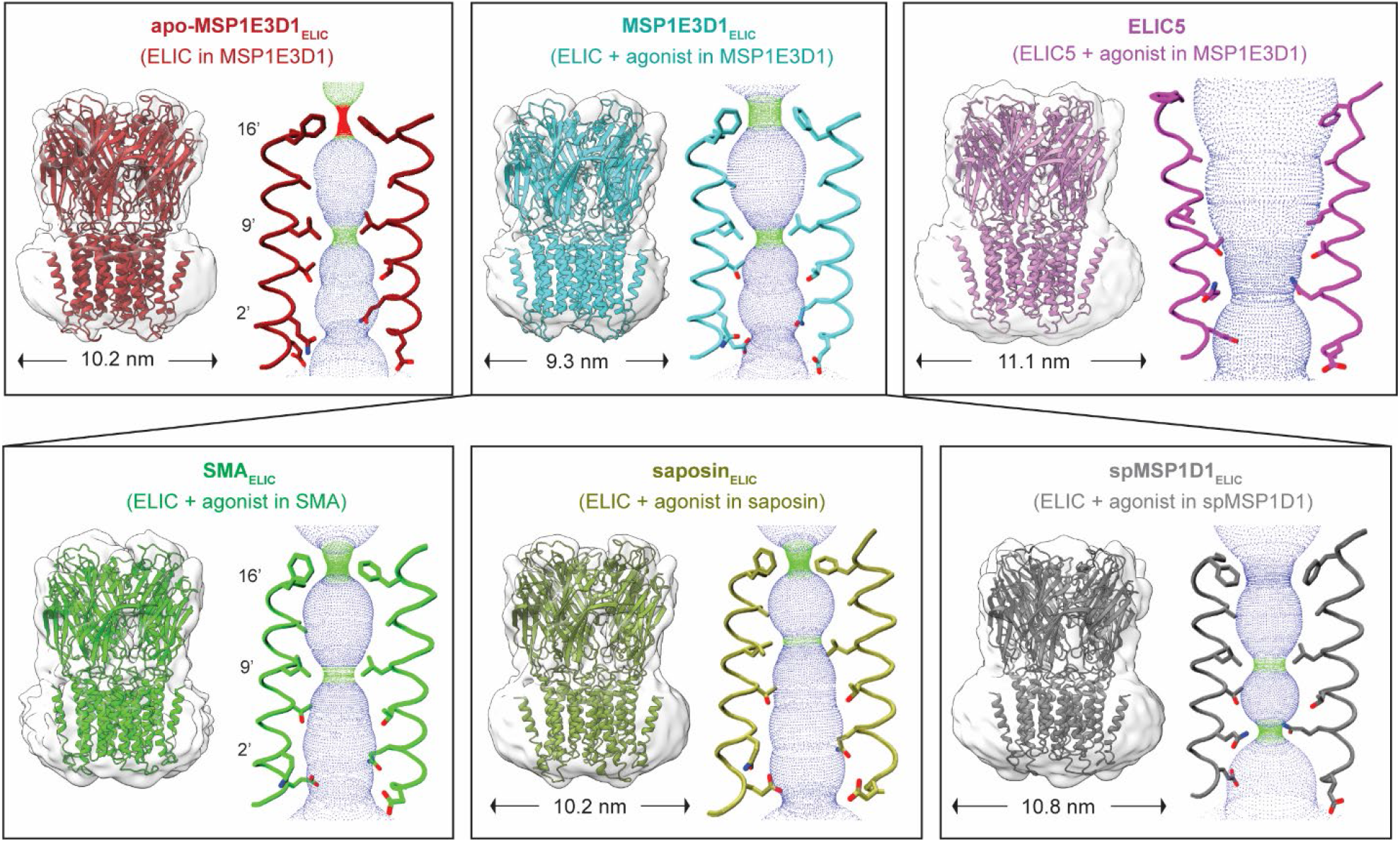
Structures and pore conformations of ELIC with different nanodisc scaffolds: Cryo-EM maps, models, and pore profile for apo-MSP1E3D1_ELIC_ (pdb 8D65), MSP1E3D1_ELIC_ (pdb 8D66), ELIC5 (pdb 8D68), SMA_ELIC_ (ELIC + 50 mM propylamine in SMA), saposin_ELIC_ (ELIC + 50 mM propylamine in saposin), and spMSP1D1_ELIC_ (ELIC + 50 mM propylamine in spMSP1D1). The measured diameter of the cryo-EM density of the nanodisc outer edge is indicated below each respective map. Cryo-EM maps were low pass filtered at 8 Å and represented in semi-transparent white. For the pore profile, two opposing M2 helices from the transmembrane domain are shown and the side chains of 16’ (F247), 9’ (L240), and 2’ (Q233) are labeled.

### spMSP1D1_ELIC_ shows constriction at the putative desensitization gate

We first examined the structure of the pore, which has been used to assign putative functional states to pLGIC structures. Apo-MSP1E3D1_ELIC_ and MSP1E3D1_ELIC_ have constrictions at the M2 pore-lining residues, 16’ (F247) and 9’ (L240), with slight opening at the top of M2 in the agonist-bound structure, consistent with putative resting and pre-active conformations (Fig. 1)^21^. The ELIC5 structure is apparently open with 2’ (Q233) forming the narrowest point of the pore (Fig. 1)^19^. Similar to MSP1E3D1_ELIC_, SMA_ELIC_ and saposin_ELIC_ show constrictions at 16’ and 9’, also consistent with pre-active conformations (Fig. 1, Supplementary Fig. 4 and 5). In contrast, spMSP1D1_ELIC_ shows a significant change in the pore architecture. Specifically, the density for the 16’ (F247) side chain is weaker suggesting a more dynamic structure; a reasonable model shows 16’ pointing intracellular partially opening the pore at this location (Fig. 1, Supplementary Fig. 4 and 5). 2’ (Q233), which is the purported desensitization gate in the muscle-type nAchR^17^ and serotonin (5-HT_3A_) receptor^22^, rotates into the pore producing another constriction with a diameter of 3.0 Å (Fig. 1, Supplementary Fig. 4 and 5). Thus, spMSP1D1_ELIC_ reveals a desensitized-like pore profile with progressive opening at the top of the pore at 16’, apparent closure at 2’, and a persistent, albeit slightly more open, constriction at 9’. This pore profile is not unlike the proposed desensitized structure of the α7 nAchR where the pore is constricted at both 9’ and the intracellular end of the pore (−1’)^23^.

Closer examination of spMSP1D1_ELIC_ in comparison with MSP1E3D1_ELIC_, SMA_ELIC_ or saposin_ELIC_ shows a series of structural changes associated with closure at 2’ (Fig. 2c-e). In MSP1E3D1_ELIC_, SMA_ELIC_ and saposin_ELIC_, 2’ (Q233) is pointing intracellular and forms intersubunit hydrogen bonds with E230 and T234 at the bottom of M2 in an adjacent subunit (Fig. 2c). However, in spMSP1D1_ELIC_, the bottom of M2 lengthens (produced by a partial unraveling of the helix), and M1 and M2 translate away from the pore (Fig. 2c-d). As a result, E230 rotates away from Q233 (2’) and now forms a hydrogen bond with S229 in the adjacent subunit. With the hydrogen bond between E230 and Q233 broken, Q233 (2’) rotates into the pore axis (Fig. 2a and 2c). Saposin_ELIC_ also shows lengthening of the bottom of M2, similar to spMSP1D1_ELIC_, such that the hydrogen bond distance between Q233 and E230 is greater at ~3.5 Å compared to ~2.5 Å in MSP1E3D1_ELIC_ and SMA_ELIC_; yet, there is less outward translation of M2 and M1 and the hydrogen bond between Q233 and T234 is maintained such that Q233 (2’) is still oriented intracellular and not into the pore (Fig. 2c, Supplementary Fig. 6a and 6b). This suggests that saposin_ELIC_ is intermediate between MSP1E3D1_ELIC_ and SMA_ELIC_, and spMSP1D1_ELIC_ with regards to this conformational change at the bottom of M2. Thus, re-arrangement of a hydrogen bond network at the bottom of M2 and M1 in ELIC leads to apparent closure of the putative desensitization gate at 2’ in spMSP1D1_ELIC_. E230 is conserved in many nAchRs and 2’ is often a serine or threonine (Supplementary Fig. 6d). Recent structures of the muscle-type nAchR also show that 2’ forms a hydrogen bond with a glutamate or glutamine in the equivalent position as E230 in ELIC^17^.

**Figure 2:**
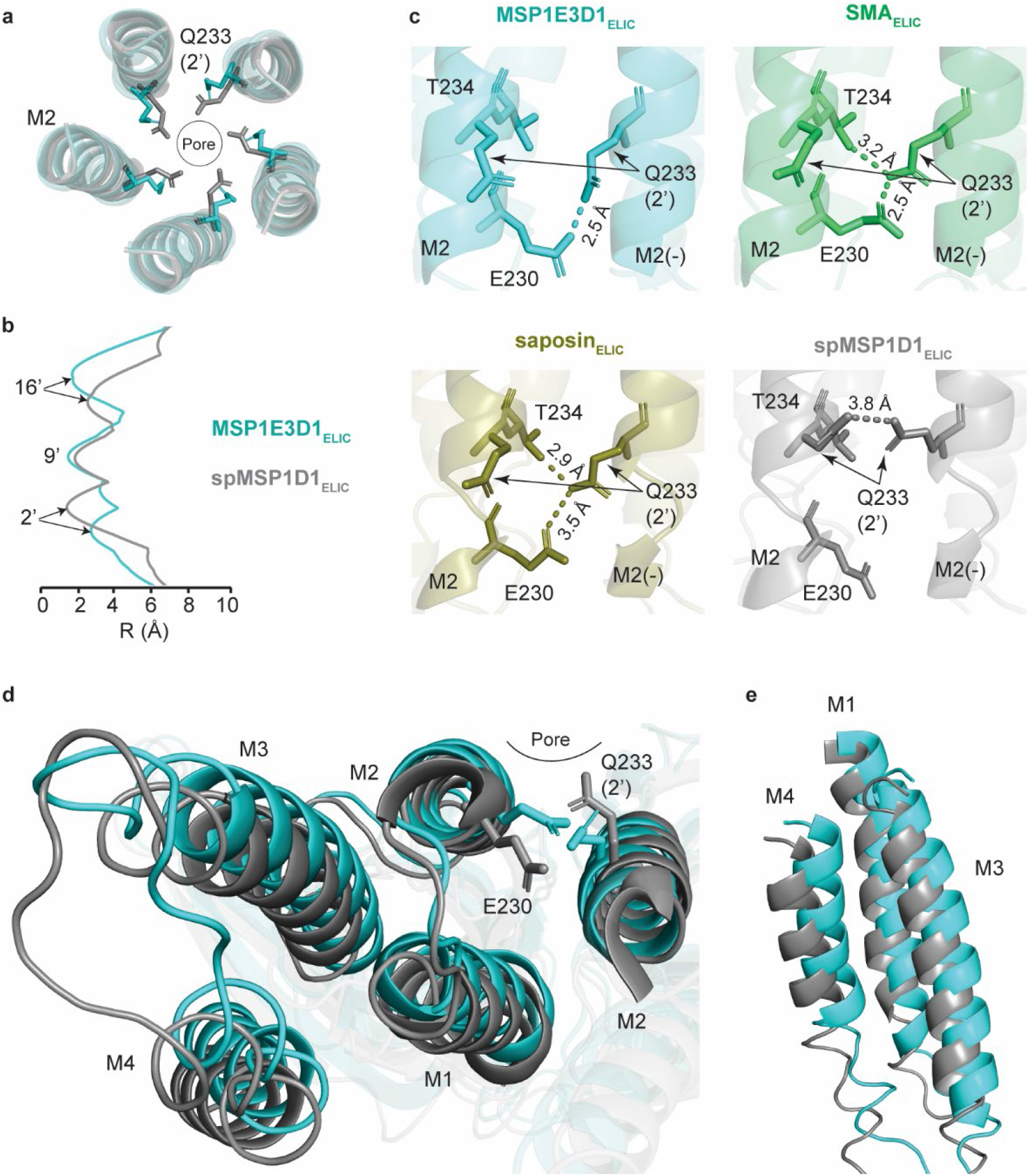
Conformational changes associated with constriction at 2’. **(a)** Top-down view of M2 and Q233 (2’) from MSP1E3D1_ELIC_ and spMSP1D1_ELIC_. The side chain of Q233 is shown. **(b)** Pore radius profile of MSP1E3D1_ELIC_ and spMSP1D1_ELIC_ measured by HOLE. **(c)** Side view of the bottom of M2 from two adjacent subunits showing an intersubunit hydrogen bond network (dashed lines) between E230, Q233, and T234. The distances between side chains is measured from the NE2 atom of Q233. Bottom-up view **(d)** and side view **(e)** of the TMD of MSP1E3D1_ELIC_ and spMSP1D1_ELIC_ showing the side chains of E320 and Q233. Images are from a global superposition of the structures.

The outward translation of M1 in spMSP1D1_ELIC_ is associated with an outward translation of M4 and M3 away from the pore (Fig. 2d and 2e). This coupled movement is evident when examining F223 and C300 at the bottom of M1 and M4, respectively, which are separated by ~4 Å and likely form an S-H/π interaction (Supplementary Fig. 6c). Interestingly, C300S or C300A mutations decrease the rate and extent of desensitization in ELIC^24,25^. This outward translation of the transmembrane helices is most pronounced in spMSP1D1_ELIC_, followed by saposin_ELIC_, and then MSP1E3D1_ELIC_ and SMA_ELIC_ (Fig. 2d and 2e, Supplementary Fig. 6a and 6b), which is consistent with the relative extent of conformational changes at the bottom of M2. Therefore, structures of ELIC in different nanodisc scaffolds reveal coupling between the packing of the transmembrane helices (M1, M3 and M4) and the putative desensitization gate at the bottom of M2. The outward translation of M4 in spMSP1D1_ELIC_ mostly closely approximates ELIC5 (fully-activated, open-channel structure), saposin_ELIC_ is intermediate, and MSP1E3D1_ELIC_ and SMA_ELIC_ are similar with the least outward translation (Supplementary Fig. 7a). Of note, the cryo-EM density of M4, especially the C-terminal end, in spMSP1D1_ELIC_, saposin_ELIC_, SMA_ELIC_ and MSP1E3D1_ELIC_ is relatively weak compared to ELIC5 (Supplementary Fig. 7b), suggesting that M4 is more dynamic in agonist-bound non-conducting conformations than in the open-channel conformation.

### Different nanodisc scaffolds produce a range of agonist-bound conformations

The structures in different nanodisc scaffolds also show significant differences in the outer transmembrane domain (TMD) and ECD (extracellular domain)-TMD interface. These differences approximately span the conformational changes associated with ELIC activation from resting to fully active, represented by apo-MSP1E3D1_ELIC_ and the ELIC5 open-channel structure, and are also similar to those reported in other pLGICs^14,17,18,22,26,27^. With activation, the top of M2 lengthens, due to partial unraveling of the helix, and tilts away from the pore. This change is most evident when comparing apo-MSP1E3D1_ELIC_ with the ELIC5 open-channel structure (Fig. 3a and 3c). Examination of the agonist-bound structures in different nanodiscs reveals incremental movement of M2 in the following order: SMA_ELIC_ ~ MSP1E3D1_ELIC_ ~ saposin_ELIC_ < spMPS1D1_ELIC_ < ELIC5. Linked to this movement of M2 is an outward (i.e. away from the pore axis) translation of the M2-M3 linker, M3 and M1 (Fig. 3d). A key residue involved in this transition is W206, which faces the membrane in the resting state and rotates into an intersubunit space displacing M3 and the M2-M3 linker with activation^19^. These conformational changes are also incrementally appreciated in each structure in the following order: apo-MPS1E3D1_ELIC_ < SMA_ELIC_ < MSP1E3D1_ELIC_ < saposin_ELIC_ < spMSP1D1_ELIC_ < ELIC5. Note that W206 faces the membrane in SMA_ELIC_ similar to apo-MPS1E3D1_ELIC_.

**Figure 3:**
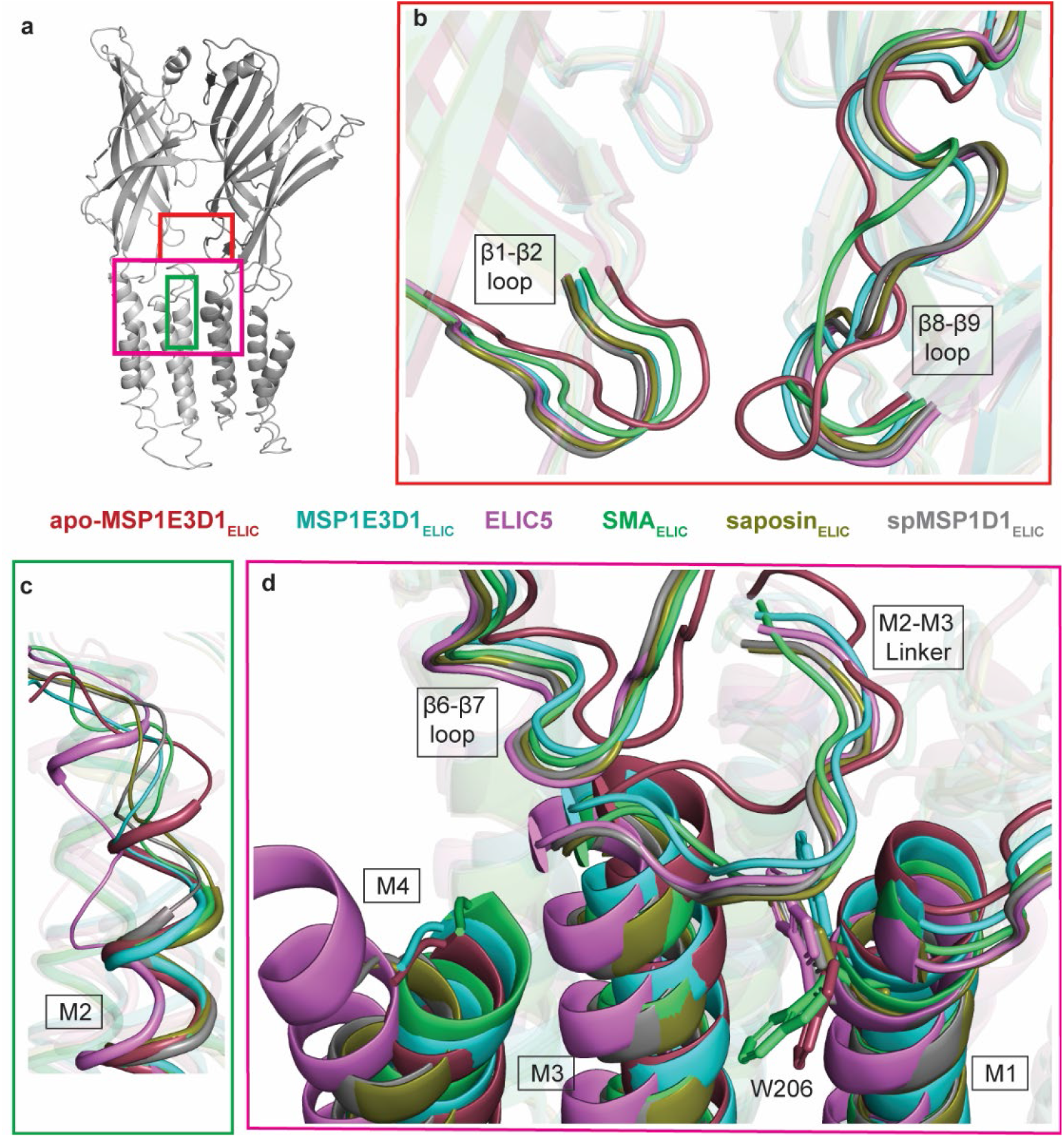
Differences in the TMD and ECD-TMD interface of ELIC in different nanodiscs scaffolds. **(a)** Two adjacent subunits of ELIC (spMSP1D1_ELIC_) with boxes indicating regions of the channel displayed in panels (b), (c) and (d). **(b)** Comparison of the β1-β2 and β8-β9 loops for the indicated structures. **(c)** Comparison of the extracellular half of M2 for the indicated structures. **(d)** Comparison of the ECD-TMD interface showing the β6-β7 loop, M2-M3 linker and the top of M1, M3 and M4 for the indicated structures. Shown is the side chain of W206. All images are a global superposition of the structures.

Inspection of the ECD-TMD loops (β6-β7, β1-β2, and β8-β9) also shows a spectrum of conformational changes in all structures associated with activation. The β6-β7 loop translates outward in the order: apo-MPS1E3D1_ELIC_ < MSP1E3D1_ELIC_ < SMA_ELIC_ < saposin_ELIC_ < spMSP1D1_ELIC_ < ELIC5 (Fig. 3d). The β1-β2 loop moves away from the adjacent subunit in the order: apo-MPS1E3D1_ELIC_ < SMA_ELIC_ < MSP1E3D1_ELIC_ ~ saposin_ELIC_ ~ spMSP1D1_ELIC_ ~ ELIC5 (Fig. 3b). The β8-β9 loop moves toward the pore vestibule in the order: apo-MPS1E3D1_ELIC_ < SMA_ELIC_ < MSP1E3D1_ELIC_ ~ saposin_ELIC_ ~ spMSP1D1_ELIC_ ~ ELIC5 (Fig. 3b). Overall, the conformational differences in the TMD and ECD-TMD interface are incremental following the trajectory of activation. With some variability, the degree of activation in agonist-bound structures is least with SMA_ELIC_, followed by MSP1E3D1_ELIC_, saposin_ELIC_, spMSP1D1_ELIC_ and ELIC5.

### Effect of nanodisc scaffold on the ECD and apo structure

Different nanodisc scaffolds also produce structural differences in the ECD. We previously noted an outward expansion of the ECD in ELIC5 compared to apo-MPS1E3D1_ELIC_ and MSP1E3D1_ELIC_, evident when comparing the β2-β3, β3-β4, β5-β6 and β7-β8 loops (Supplementary Fig. 8)^19^. Interestingly, SMA_ELIC_, saposin_ELIC_ and spMSP1D1_ELIC_ all resemble ELIC5 with respect to the more expanded ECD (Supplementary Fig. 8). To determine if this expanded ECD is a function of agonist, we determined a structure of ELIC in spMSP1D1 in the absence of agonist (Supplementary Table 1, Supplementary Fig. 1).

We call this structure apo-spMS1PD1_ELIC_. Strikingly, the ECD of apo-spMS1PD1_ELIC_ is also expanded similar to SMA_ELIC_, saposin_ELIC_ and spMSP1D1_ELIC_ (Supplementary Fig. 8). We also examined the propylamine-occupied agonist binding site. In apo-spMSP1D1_ELIC_, loop C, which caps the agonist binding site in the ECD, is oriented outward and the size of the agonist binding site is largest (Fig. 4a-b). In the agonist-bound structures, loops C moves inward toward the pore vestibule making the agonist binding site smaller in the following order from largest to smallest: SMA_ELIC_ > saposin_ELIC_ > spMSP1D1_ELIC_ > ELIC5 (Fig. 4a-c). Thus, movement of loop C and the size of the agonist binding site in SMA_ELIC_, saposin_ELIC_, and spMSP1D1_ELIC_ correlates with the degree of conformational change in the TMD and ECD-TMD interface associated with activation (i.e. structures with greater activation show a smaller agonist binding site). This correlation is consistent with the structural effect of full and partial agonists in the glycine receptor^14^.

**Figure 4:**
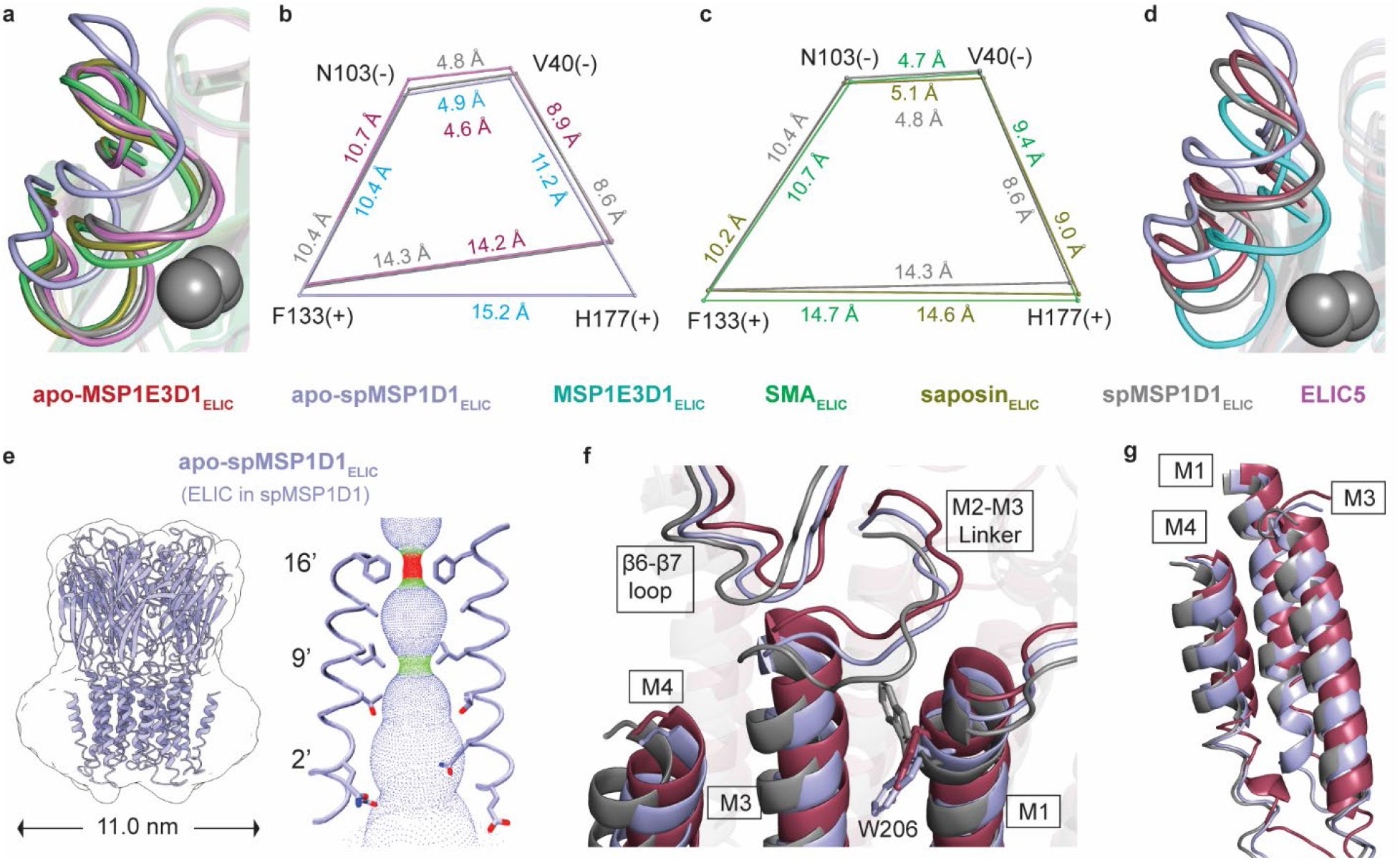
Impact of nanodisc scaffold on the apo structure and agonist-binding site. **(a)** Comparison of Loop C in apo-spMSP1D1_ELIC_, SMA_ELIC_, saposin_ELIC_, spMSP1D1_ELIC_, and ELIC5. For clarity, only propylamine from spMSP1D1_ELIC_ is shown in gray spheres. Images are from a global superposition of the structures. **(c)** and **(d)** Distances (Å) between four amino acid residues (F133(+), H177(+), V40(−), and N103(−)) in the agonist binding site comparing the indicated structures. Residue points and line distances are drawn to scale from global superposition of the structures. **(d)** Same as (a) comparing apo-MSP1E3D1_ELIC_, MSP1E3D1_ELIC_, apo-spMSP1D1_ELIC_, and spMSP1D1_ELIC_. **(e)** Cryo-EM map, model, pore profile of apo-spMSP1D1_ELIC_ (ELIC without agonist in spMSP1D1) as in Figure 1. The measured diameter of the cryo-EM density of nanodisc scaffolds is indicated below the map. **(f)** Comparison of the ECD-TMD interface showing the β6-β7 loop, M2-M3 linker and the top of M1, M3 and M4 for the indicated structures. The side chain of W206 is shown. **(g)** Side view of the TMD of the indicated structures. All images are a global superposition of the structures.

A a counter-clockwise twisting of the ECD, when viewed from the extracellular space, can be appreciated in spMSP1D1_ELIC_, saposin_ELIC_, and SMA_ELIC_ relative to apo-spMSP1D1_ELIC_ (Supplementary Fig. 9a), which is a conserved conformational change associated with activation in all pLGICs^18,22,26,28,29,30,31,32^. In contrast, MSP1E3D1_ELIC_ shows minimal counter-clockwise twisting of the ECD compared to apo-MSP1E3D1_ELIC_ (Supplementary Fig. 9b). Moreover, while MSP1E3D1_ELIC_ shows inward movement of loop C and capping of the agonist binding site relative to apo-MSP1E3D1_ELIC_, the agonist binding site is smaller in both MPS1E3D1 structures compared to the respective apo and agonist-bound spMSP1D1 structures (Fig. 4d). The TMD is also significantly different between apo-spMSP1D1_ELIC_ and apo-MSP1E3D1_ELIC_. While the pore profile of apo-spMSP1D1_ELIC_ is consistent with a resting conformation (Fig. 4e), M1, M3 and M4, along with the M2-M3 linker and β6-β7 loop, are translated outward in apo-spMSP1D1_ELIC_ compared to apo-MSP1E3D1_ELIC_ (Fig. 4f and 4g, Supplementary Fig. 10a). Relative to apo-spMSP1D1_ELIC_, spMSP1D1_ELIC_ shows an even greater outward translation of these structures characteristic of activation in the TMD (Fig. 4f and 4g). In contrast, MSP1E3D1_ELIC_ shows no outward translation or expansion of the TMD helices compared to apo-MSP1E3D1_ELIC_, indicating that this scaffold significantly limits channel activation (Supplementary Fig. 10b). In summary, the MSP1E3D1 scaffold uniquely produces a more compact ECD and TMD in both apo and agonist-bound structures with limited activation in the presence of agonist.

Having obtained apo-spMSP1D1_ELIC_, we observed a notable pattern in the conformation of M4 when comparing all structures. We added the previously-determined apo structure of ELIC in SMA (apo-SMA_ELIC_, PDB 7L6Q) to the comparison^33^. The orientation of M4 suggests two distinct groups of ELIC structures. If we associate apo-spMSP1D1_ELIC_ and ELIC5 with resting and activated conformations, spMSP1D1_ELIC_ and saposin_ELIC_ show intermediate conformations of M4 along the apparent trajectory of this helix (Fig. 5). In contrast, apo and agonist-bound structures in MSP1E3D1 and SMA show an M4 that deviates from this trajectory, packed closer to the M1 and M3 helices, with minimal change between apo and agonist-bound structures (Fig. 5). Therefore, the orientation of M4 indicates that MSP1E3D1_ELIC_ and SMA_ELIC_ are distinct from spMSP1D1_ELIC_ and saposin_ELIC_; the former show a more compact TMD and limited change with agonist, while the latter appear to be intermediates between apo-spMSP1D1_ELIC_ and ELIC5.

**Figure 5:**
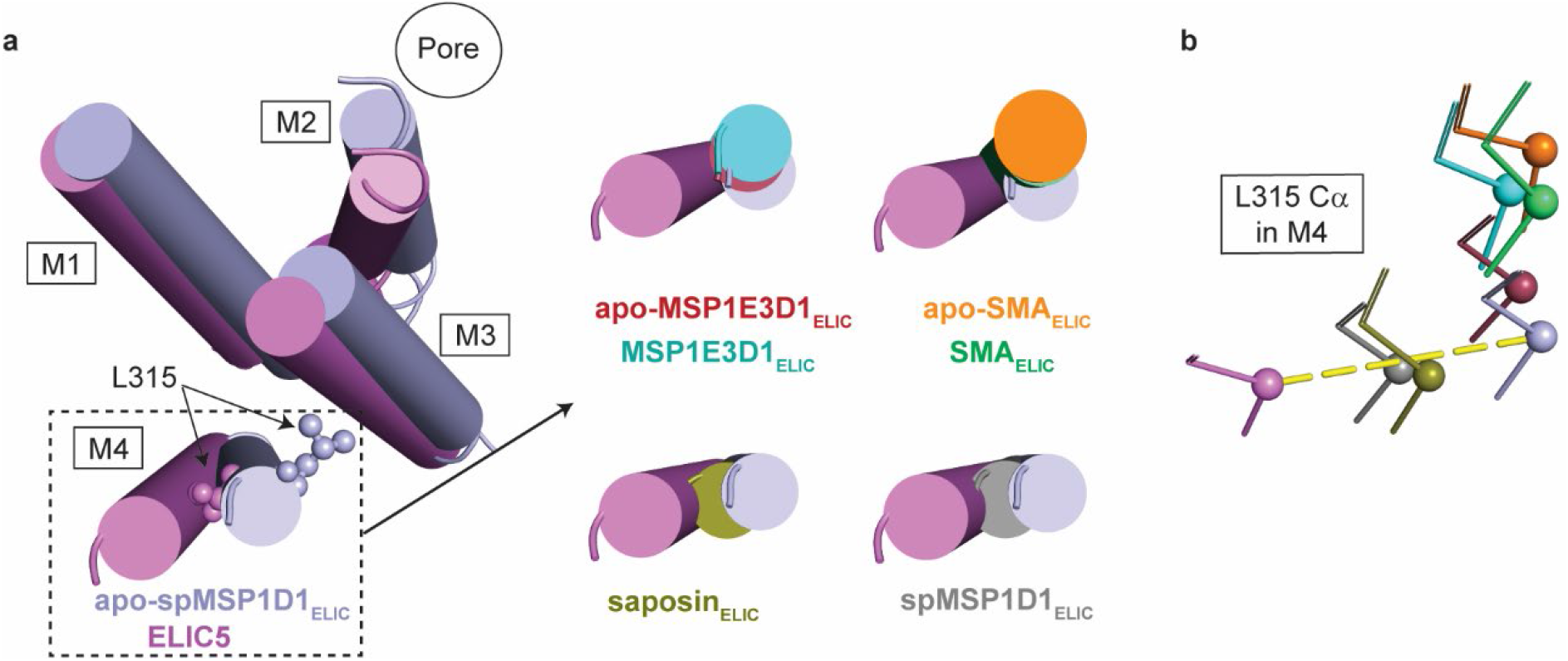
Comparison of the orientation of M4. **(a)** Top-down view of the TMD helices comparing apo-spMSP1D1_ELIC_ and ELIC5, along with four other top-down views of M4 comparing the indicated structures with apo-spMSP1D1_ELIC_ and ELIC5. Apo-SMA_ELIC_ is PDB 7L6Q (https://www.wwpdb.org/pdb?id=pdb_00007L6Q). The M4 helix closely overlaps in apo-MSP1E3D1_ELIC_ and MSP1E3D1_ELIC_, and apo-SMA_ELIC_ and SMA_ELIC_. **(b)** Comparison of the peptide backbone atoms of L315 in M4 of all structures from the same perspective as in (a) (Cα atoms in spheres). A dashed yellow line is placed connecting apo-spMSP1D1 and ELIC5. All images are taken from a global superposition of the structures.

### MD simulations of ELIC in nanodiscs

In this study, structures of ELIC in different nanodisc scaffolds produce distinct conformations via cryo-EM. What is responsible for these conformational differences, however, is unclear. Nanodiscs perturb multiple lipid physical properties^6,7,9^, and pLGICs have been shown to be sensitive to membrane thickness^34^. In addition, it is possible that interactions between the MSP (membrane scaffold protein) and ELIC could be responsible for altering the dynamics of the TMD. There is a correlation between the estimated size of the nanodisc and the degree of activation in the agonist-bound ELIC structures, with larger diameter nanodiscs demonstrating conformations that are more activated (i.e. show similarity to ELIC5). MSP1E3D1_ELIC_ and spMP1D1_ELIC_, both reconstituted in a MSP-based scaffold, produced ~9-10 and ~11 nm nanodisc particles, respectively. Thus, we explored the influence of nanodisc size on ELIC structure by performing 500 ns of molecular dynamics (MD) simulations of spMP1D1_ELIC_ in MSP-based 9 and 11 nm nanodiscs (Fig. 6a, Supplementary Fig. 11). Since nanodiscs are thought to recapitulate a lipid bilayer environment, we also analyzed a simulation of spMP1D1_ELIC_ in a planar bilayer. By using the same starting structure (spMSP1D1_ELIC_) in the simulations, we can directly examine the effect of nanodisc size on ELIC structure.

**Figure 6:**
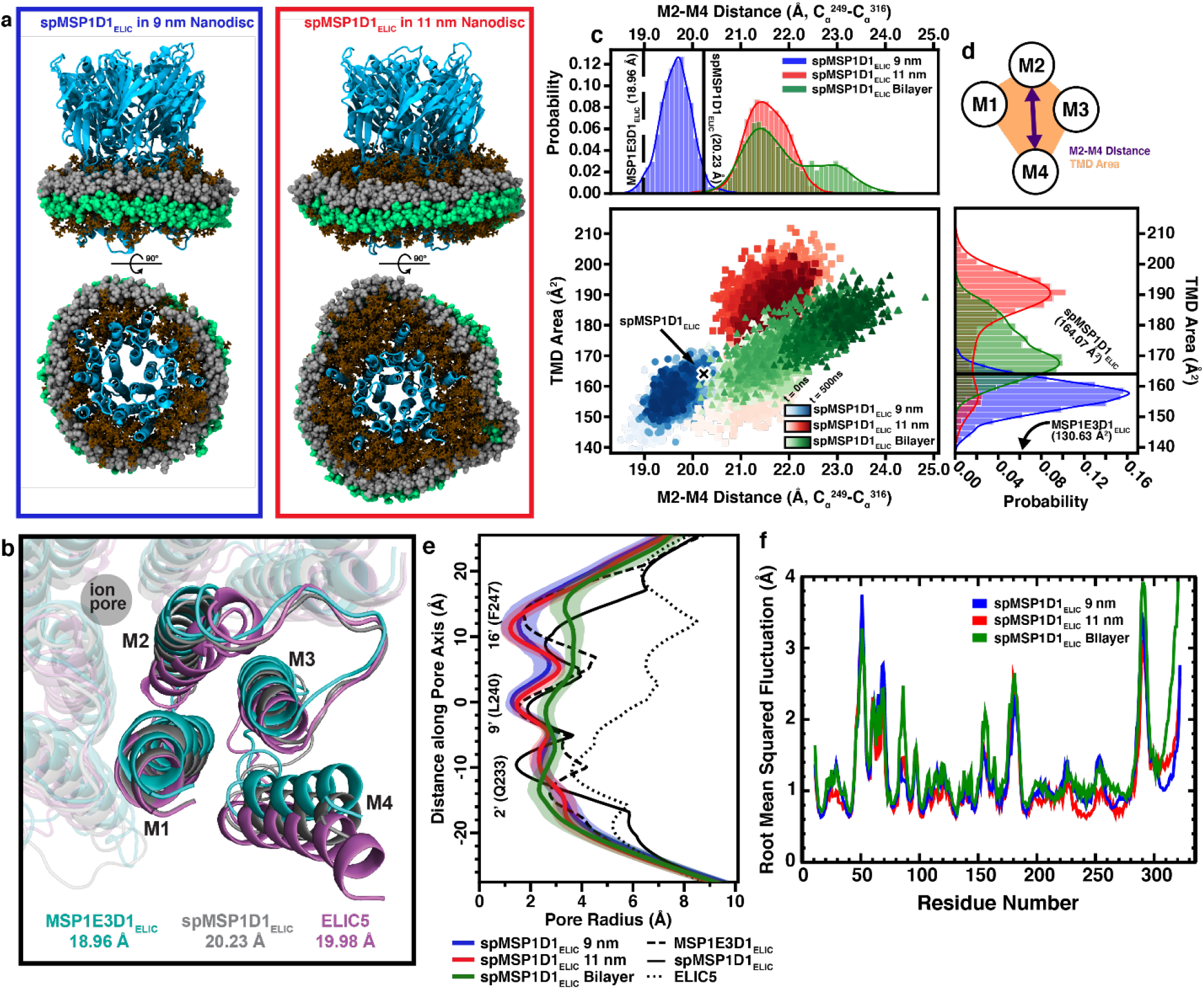
Structure and dynamics of ELIC in a 9 nm versus an 11 nm nanodisc. **(a)** Atomistic models of spMSP1D1_ELIC_ equilibrated in a 9 nm (left) and 11 nm (right) nanodisc. spMSP1D1_ELIC_ is cyan, MSP silver and green, and POPC brown licorice. For the top-down view, the ECD was removed to improve clarity of the TMD and MSP. **(b)** Cryo-EM structures of the MSP1E3D1_ELIC_, spMSP1D1_ELIC_, and ELIC5 TMD superimposed globally to demonstrate the difference in M2-M4 distance (shown under each structure label). **(c)** Two-dimensional plot of M2-M4 distance versus TMD area as a function of simulation time for the 9 nm nanodisc model (spMSP1D1_ELIC_ 9 nm, blue), 11 nm nanodisc model (spMSP1D1_ELIC_ 11 nm, red), and lipid bilayer model (spMSP1D1_ELIC_ bilayer, green). In the scatter plot, darker shading indicates later time points in the simulation. The starting point for the simulation is denoted by “X” and labeled “spMSP1D1_ELIC_”. Histograms showing the probability for M2-M4 distance (top) and TMD area (right) are shown. **(d)** A cartoon representation of how the M2-M4 distance and TMD area are measured. **(e)** Pore radius profile in the spMSP1D1_ELIC_ 9 nm, spMSP1D1_ELIC_ 11 nm, and spMSP1D1_ELIC_ bilayer simulations. The heavy trace represents the average over the last 250 ns of the simulation and the transparent area +/- 1 standard deviation. The position of the 2’ (Q233), 9’ (L240), and 16’ (F247) side chains are shown, as well as the pore radius profiles for MSP1E3D1_ELIC_, spMSP1D1_ELIC_, and ELIC5. **(f)** Root mean squared fluctuation (RMSF) of ELIC for the spMSP1D1_ELIC_ 9 nm, spMSP1D1_ELIC_ 11 nm, and spMSP1D1_ELIC_ bilayer simulations. The RMSF was calculated using the Cα of each residue in ELIC and averaged over the last 250 ns of each simulation.

There is a global outward blooming of the transmembrane helices M1, M3 and M4, when comparing the fully-activated ELIC5 structure with all other structures. This is evidenced by an increase in the distance between M2 and M4, and the area formed by all four transmembrane helices (Fig. 6b). Similarly, the M2-M4 distance and TMD area are larger in spMP1D1_ELIC_ compared to MSP1E3D1_ELIC_ (Fig. 6b). Using T249 and V316 to represent the position of M2 and M4, respectively, we examined the M2-M4 distance and TMD area (with M1 represented by I209 and M3 represented by I262) in our simulations (Fig. 6c-d). There is a clear difference in these measurements between the two nanodisc systems. In the 9 nm nanodisc, we observed a contraction of the M2-M4 distance and the overall TMD area from the starting spMP1D1_ELIC_ structure, suggesting that the smaller nanodisc is forcing compaction of the TMD. In contrast, the larger, 11 nm nanodisc showed a greater M2-M4 distance and TMD area, consistent with the difference between spMP1D1_ELIC_ and MSP1E3D1_ELIC_. Interestingly, the bilayer simulation shows a metastable conformation that is similar to the 11 nm nanodisc model with respect to the M2-M4 distance; however, M4 continues to translate even further away from M2 later in the bilayer simulation. The TMD area in the bilayer simulation is intermediate between the 9 and 11 nm nanodisc simulations. We also compared the C_α_ root-mean-square fluctuation (RMSF) of ELIC (Fig. 6f). Overall, the dynamics of the ELIC TMD are constrained in the nanodisc systems compared to the planar bilayer, as evidenced by lower RMSF in the nanodisc systems. Of note, M4, especially residues 305-322 shown to be important in desensitization of the channel, is very dynamic in the planar bilayer simulation compared to the nanodisc systems (Fig. 6f)^35^. However, M4 fluctuations are still higher in the 11 nm nanodisc than 9 nm nanodisc. These results suggest that the 11 nm nanodisc represents an environment that is closer to a planar bilayer than the 9 nm nanodisc, but that neither nanodisc system supports the same dynamics as a planar bilayer. Indeed, when we examine the pore radius profile of these three simulation systems (Fig. 6e), we note that the shape of the profile in the planar bilayer is distinct from that of both nanodisc simulation systems, with both the 9 and 11 nm nanodisc systems showing a pre-active or resting-like profile (constriction at the 9’ and 16’ positions and a relative opening at the 2’ position compared to the starting structure, spMSP1D1_ELIC_) while the same structure in a planar bilayer shows a profile that is more consistent with an activated or desensitized-like conformation (relative widening of the 9’ and 16’ positions). We conclude that the nanodisc, including nanodisc size, has a profound effect on ELIC TMD structure and dynamics.

Two possible explanations for the difference in ELIC TMD behavior observed in our nanodisc systems include alterations to the bilayer physical properties and direct interactions between ELIC and the MSP scaffold. Empty nanodiscs are reported to have position-dependent changes in membrane thickness, being relatively thin at the nanodisc rim to minimize hydrophobic exposure and relatively thick in the nanodisc center^6,7^. We first measured the bilayer thickness as a function of distance from the ELIC surface. In the 9 nm nanodisc, both the overall membrane thickness and the thickness of the hydrophobic core are significantly reduced compared to the 11 nm nanodisc and the planar bilayer, which show similar membrane thickness profiles (Fig. 7a). Additionally, we examined the two-dimensional membrane thickness for all three simulation systems. There is a localized elevation of the membrane surrounding the M4 helix that is present in both the bilayer and 11 nm nanodisc that is not present in the 9 nm nanodisc simulation (Supplemental Fig. 12).

**Figure 7:**
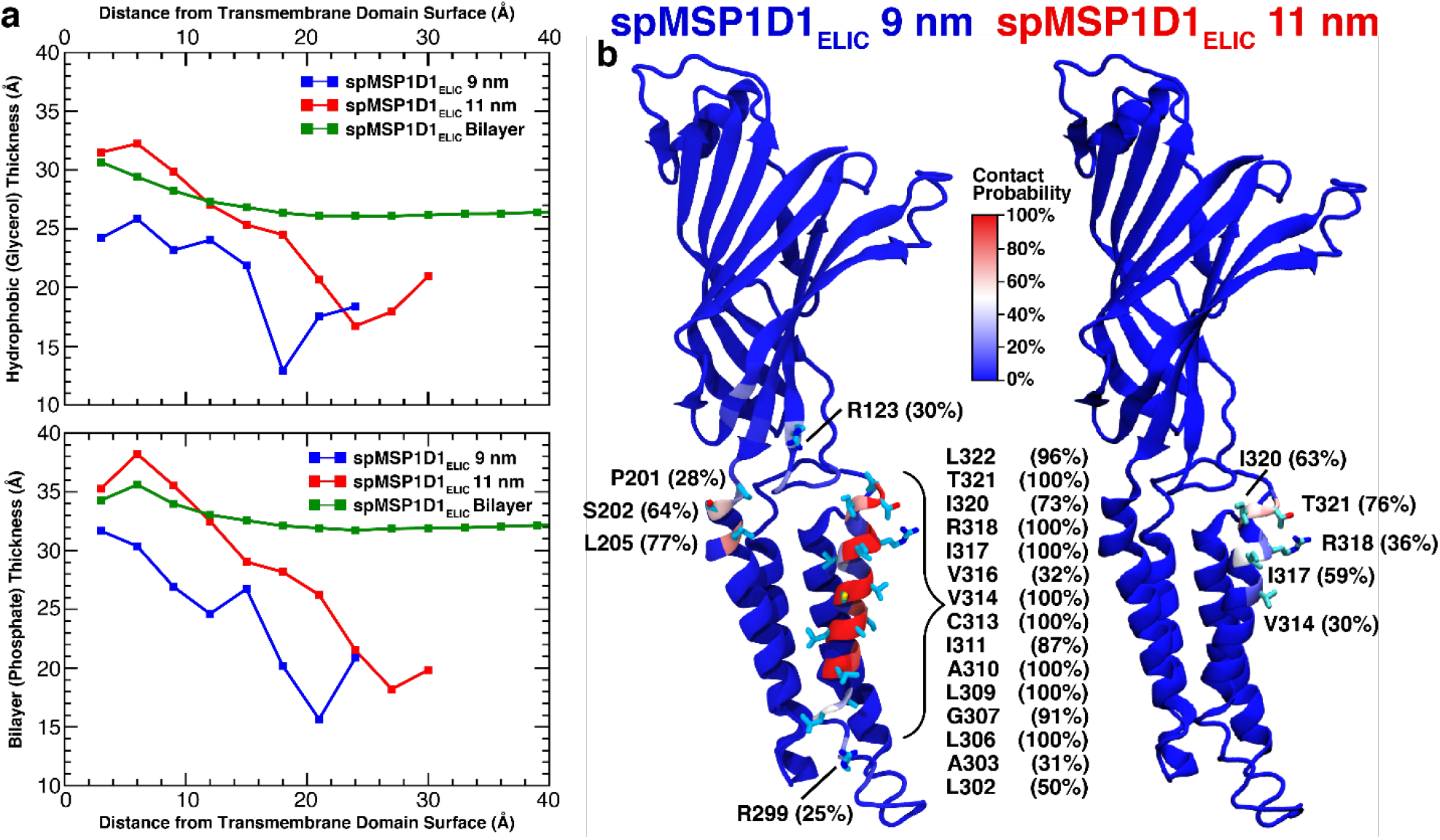
Quantifying the effect of the nanodisc environment. **(a)** The hydrophobic and total bilayer thickness are displayed relative to distance from the ELIC TMD surface for the spMSP1D1_ELIC_ 9 nm, spMSP1D1_ELIC_ 11 nm, and spMSP1D1_ELIC_ bilayer simulations. Data for the bilayer simulation more than 40 Å away from the protein surface was excluded for clarity. **(b)** The probability of ELIC contacting the MSP is shown for the spMSP1D1_ELIC_ 9 nm and spMSP1D1_ELIC_ 11 nm simulations. The probability was calculated over the last 250 ns of the simulation with a contact denoted by any atom of a given ELIC residue within 4.5 Å of any atom in the MSP. Residues with low contact probability are shown in blue and high contact probability is shown in red. Residues with a >25% contact probability are explicitly shown as licorice representations with the calculated contact probability displayed. Most resides with high contact probability are in M4.

In addition to changes in membrane thickness, we also observed significant direct interaction between ELIC and the MSP (Fig. 7b). In the 9 nm nanodisc simulation, there is constant interaction between the outer face of M4 and the MSP as well as some interaction between the MSP and the β6-β7 loop. While there are M4-MSP interactions in the 11 nm nanodisc simulation, the number of interaction sites and the frequency of interaction is significantly reduced compared to the 9 nm nanodisc system. In the 11 nm simulation, only interaction between the MSP and the C-terminal residues of the M4 helix are observed. Therefore, direct interactions with the membrane scaffold could also be altering the conformational dynamics of ELIC, as evidenced by the reduced C_α_ RMSF of M4 in the nanodisc simulations (Fig. 6f).

## Discussion

The agonist-bound structures of ELIC in different nanodiscs can be categorized into two groups: 1) those with limited activation (MSP1E3D1 and SMA), and 2) those which show significant activation (saposin and spMSP1D1). MSP1E3D1 and SMA produce structures with limited conformational changes between apo and agonist-bound structures, especially in the TMD. In the MSP1E3D1 structures, the ECD is uniquely contracted and the counter-clockwise twisting of the ECD that is characteristic of pLGIC activation is absent. In both MSP1E3D1_ELIC_ and SMA_ELIC_, the TMD is contracted with minimal outward movement of the M1, M3 and M4 helices compared to the apo structures. Moreover, the TMD of apo-MSP1E3D1_ELIC_ and apo-SMA_ELIC_ (PDB 7L6Q) are more compact than apo-spMSP1D1_ELIC_. Therefore, a general pattern in these structures is that nanodiscs which limit activation produce a more compact TMD in both apo and agonist-bound structures.

In contrast, saposin_ELIC_ and spMSP1D1_ELIC_ show an outward expansion of the TMD (M1, M3 and M4) compared to apo-spMSP1D1_ELIC_ that mimics the putative, activated ELIC5 structure. This is especially apparent when comparing the orientation of M4. In saposin_ELIC_ and spMSP1D1_ELIC_, M4 follows the apparent trajectory of this helix from apo-spMSP1D1_ELIC_ to ELIC5. Saposin_ELIC_ and spMSP1D1_ELIC_ also show concerted conformational changes in the ECD and ECD-TMD interface like ELIC5 only to a lesser degree. Thus, if one considers apo-spMSP1D1_ELIC_ and ELIC5 to be resting and activated conformations, respectively, saposin_ELIC_ and spMSP1D1_ELIC_ appear to be intermediate conformations. The pore profiles of saposin_ELIC_ and spMSP1D1_ELIC_ suggest that they are pre-active and desensitized-like conformations, respectively. However, while spMSP1D1_ELIC_ shows a constriction at 2’, this is lost in the MD simulations even in a lipid bilayer. Therefore, the MD simulations of spMSP1D1_ELIC_ suggest that this structure is not very stable, making functional annotation uncertain. Further work is needed to annotate these structures and to identify stable, desensitized conformations of ELIC. We previously noted an apparent discrepancy between functional measurements of ELIC in 2:1:1 membranes, which indicate that agonist-bound ELIC should be desensitized, and the structure of agonist-bound ELIC in 2:1:1 MSP1E3D1 nanodiscs, which shows an apparent pre-active conformation^19^. While the functional annotation of the ELIC structures remains a challenge, the results of this study indicate that this difference is likely a consequence of the nanodisc limiting activation of the channel.

The profound effect of nanodiscs on ELIC structure raises the unsettling question of whether any of these structures are sampled in a native membrane environment. This question may not be definitively answered without structures of ELIC in a lipid bilayer such as a liposome. However, it is noteworthy that the differences in the agonist-bound structures are generally concerted, falling within a spectrum of conformational changes associated with channel activation ^19^ in an order of least to most activated of: SMA < MSP1E3D1 < saposin < spMSP1D1. More activated structures produce greater capping of the agonist binding site, and more outward movement of the β6-β7 loop, M2-M3 linker and the M1, M3 and M4 helices. Broadly, these agonist-induced conformational changes agree with those reported in other pLGICs^14,17,18,22,26,27^. Strikingly, different nanodisc scaffolds produce changes in the ECD and agonist binding site, indicating that the nanodisc environment can support long-range conformational changes in pLGIC structure. These regions, such as the agonist binding site, are ~40 Å away from the TMD and lipids rendering direct effects of the nanodisc unlikely. Moreover, in spMSP1D1_ELIC_, the lengthening of M2 and outward translation of the transmembrane helices leads to re-arrangement of a hydrogen bond network at the intracellular end of M2 and constriction at 2’. Therefore, the structures of ELIC in different nanodiscs demonstrate features of allostery in this ion channel. The nanodisc scaffold dictates whether the TMD expands with agonist binding. With the exception of MSP1E3D1 structures, a more expanded TMD is associated with a more contracted agonist binding site. Expansion of the TMD also produces a lengthening of M2 and constriction at the intracellular end of the pore.

The effect of nanodisc scaffold on ELIC activation correlates with nanodisc diameter: larger nanodiscs (saposin or spMSP1D1) produce a more activated structure with an expanded TMD. MD simulations of spMSP1D1_ELIC_ in a 9 nm nanodisc, which approximates MSP1E3D1_ELIC_, and 11 nm nanodisc, which approximates spMSP1D1_ELIC_, produce differences in the M2-M4 distance and TMD area that follow the same trend as in the structures (i.e. MSP1E3D1_ELIC_ and spMSP1D1_ELIC_) suggesting the importance of nanodisc size. Interestingly, the RMSF of M4 in the 11 nm nanodisc model are more similar to the planar lipid bilayer relative to the 9 nm nanodisc model. This may explain why the larger circularized nanodisc (spMSPD1) supports a more expanded TMD, and seemingly activated channel. Still, the structure and dynamics of M4 in the 11 nm nanodisc model do not match the planar lipid bilayer model suggesting that this larger nanodisc system does not fully mimic the bilayer. Furthermore, in both the 9 and 11 nm nanodisc simulations, the pore structure of ELIC, which starts with a constricted 2’ and relatively open 16’, changes to a pre-active or resting-like conformation (i.e. constriction at 16’ and widening at 2’). In contrast, the planar bilayer simulation shows a more activated conformation (i.e. further opening at 16’ and 9’). This suggests that even the 11 nm nanodisc system alters the structure of ELIC compared to a planar bilayer, seemingly restricting gating. The results indicate that nanodisc size impacts ELIC structure, and suggests the possibility that larger circularized nanodiscs may provide better membrane mimetics.

How the nanodisc alters ELIC structure is a challenging question to address. The simulations suggest possible factors such as membrane thickness and curvature, or direct interactions between the scaffold and M4. It is intriguing that the scaffold makes frequent contacts with M4 in the 9 nm nanodisc model, and this is associated with a restriction of M4 dynamics and transmembrane helices that are more tightly packed. This effect is likely complicated by differences in the structure or chemistry of the scaffold that the models in the MD simulations do not account for. Nevertheless, it is noteworthy that the diameter of the cryo-EM density of the GABA_A_R in MSP2N2 nanodiscs was also estimated to be ~9 nm^15^; thus, the GABA_A_R in conventional MSP nanodiscs is likely to also interact with the nanodisc scaffold, especially through M4, possibly altering channel structure. This may explain why a recent structure of the α1β3 GABA_A_R showed a closed activation gate despite having an estimated peak open probability of ~0.6 in HEK293 cells^30^. It may also explain why systematic differences were noted between α1β3γ2 and α1β2γ2 GABA_A_R structures in MSP2N2 and saposin nanodiscs, respectively^15,36^. Structures in the presence of GABA and picrotoxin showed greater activation in saposin than in MSP2N2 (i.e. a wider pore and more expanded TMD in the saposin structure), similar to our finding that saposin produces a more activated structure of ELIC than MSP1E3D1. In another study, a structure of agonist-bound GlyR in MSP2N2 produced a desensitized conformation, while agonist-bound GlyR in SMA produced a mixture of pre-active, open-channel and desensitized conformations^14^. While we have not identified multiple conformations of ELIC from a single dataset, we also find that SMA limits activation of ELIC. Therefore, the effect of different nanodisc scaffolds on ELIC may be present to varying degrees in other pLGICs such as the GABA_A_R and GlyR.

In conclusion, the nanodisc scaffold affects the structure of ELIC and this effect likely relates, in part, to nanodisc size. Smaller nanodiscs restrict ELIC activation, which may be a consequence of direct interactions between the scaffold and M4. A larger circularized nanodisc, spMSP1D1, produced an ELIC structure most similar to the fully-activated ELIC5 structure. This is similar to a structural study of TRPV3, which used a circularized nanodisc to capture activation by heat^37^. It will be critical to consider the effect of nanodisc scaffolds when studying the structure of pLGICs and possibly other membrane proteins. One perspective is that certain nanodiscs are better membrane mimetics and more likely to produce functionally-relevant conformations. Another is that multiple nanodiscs can be employed to better sample the conformational landscape of a membrane protein.

## Methods

### Purification and reconstitution of ELIC in nanodiscs

ELIC was expressed and purified as previously described^38^, using pET-26-MBP-ELIC provided by Raimund Dutzler (Addgene plasmid #39239). ELIC was expressed in OverExpress C43 (DE3) *E. coli* (Lucigen) with Terrific Broth (Sigma) using 0.1 mM IPTG for induction. The cells were lysed using an Avestin C5 emulsifier, isolated membranes were solubilized with 1% DDM (Anatrace) and purified using amylose resin (New England Biolabs). The protein was eluted with buffer A (10 mM Tris pH 7.5, 100 mM NaCl) plus 0.02% DDM, 0.05 mM TCEP (ThermoFisher Scientific) and 40 mM maltose (Sigma), digested overnight with HRV-3c protease (ThermoFisher Scientific), and purified over a Sephadex 200 Increase 10/300 size exclusion column (Cytiva) in buffer A with 0.02% DDM.

Reconstitution of ELIC in saposin and spMSP1D1 was performed using a liposome destabilization technique^19^. A 2:1:1 molar mixture of POPC:POPE:POPG in chloroform was dried overnight in a desiccator, rehydrated in buffer A to 7.5 mg/ml (~10 mM), freeze-thawed 3x and extruded with a 400 nm filter (Avanti Polar Lipids). These liposomes were destabilized with DDM at a final concentration of ~0.4% DDM at RT for 3 h. Next, ELIC and the nanodisc scaffold protein was added at the following ELIC:scaffold:phospholipid molar ratios: 1:30:230 for saposin, and 1:2:290 for spMSP1D1. This mixture yielded a final DDM concentration of ~0.2% and was rotated RT for 1.5 h, followed by Bio-beads SM-2 Resin (Bio-Rad) for the removal of DDM overnight at 4 °C. The nanodisc sample was finally purified over a Sephadex 200 Increase 10/300 column in 10 mM HEPES pH 7.5 with 100 mM NaCl. Propylamine was added to a final concentration of 50 mM at least 30 min prior to freezing for cryo-EM. For saposin and spMSP1D1, samples were concentrated to ~1.2 mg/ml. The His-tagged saposin construct was obtained from Salipro Biotech AB, and purified using a Ni-NTA column. After removal of the His-tag using TEV digestion, the protein was purified by size exclusion chromatography. spMSP1D1 was a gift from Huan Bao (Addgene plasmid #173482), and purified using Ni-NTA without size exclusion chromatography^5^.

Reconstitution of ELIC in SMA was performed by first generating ELIC proteoliposomes. 2:1:1 POPC:POPE:POPG was solubilized in buffer A with ~40 mM CHAPS, after which ELIC was added (100 μg per mg of lipid) for 30 min at RT. Next BioBeads was added to remove DDM rotating for ~2.5 hr. The proteliposome suspension was extruded with a 100 nm filter (Avanti Polar Lipids). To form SMA nanodiscs, 20% SMALPs 300 (Orbiscope) was added to the Biobead-free proteoliposomes to a final concentration of 2.5%, and agitated at RT for 2 hours in the absence of light. To isolate ELIC SMA nanodiscs from empty SMA nanodiscs, the sample was purified over a Ni-NTA column (WT ELIC binds to Ni-NTA likely through native histidine residues) and eluted with 30 mM imidazole. The eluate was then purified over a Sephadex 200 Increase 10/300 column in 10 mM HEPES pH 7.5 with 100 mM NaCl. Fractions containing the nanodisc sample were concentrated to ~1.2 mg/ml and propylamine was added to 50 mM for cryo-EM imaging. **Cryo-EM sample preparation and imaging**. 3 μl of ELIC nanodiscs at ~1.2 mg/ml were pipetted onto Quantifoil R2/2 copper grids (which had previously been cleaned using a Gatan Solaris 950 in a H_2_/O_2_ plasma for 60 s), after which each grid was blotted for 2 s in a 100% humidity environment and vitrified in liquid ethane using a Vitrobot Mark IV (ThermoFisher Scientific). Each grid was then imaged on a Titan Krios 300 kV Cryo-EM equipped with a Falcon 4 Direct Electron Detector (ThermoFisher Scientific). Single particle cryo-EM data was acquired using counting mode on the Falcon 4 with the EPU software (version 2.12.1.2782). Movies were collected using a pixel size of 0.9 Å for SMA_ELIC_ and 0.7 Å for saposin_ELIC_, spMSP1D1_ELIC_ and apo spMSP1D1_ELIC,_ with a defocus range of −0.8 to −2.4 μm. For SMA_ELIC_, each movie consisted of 46 individual frames with a per-frame exposure time of 200 ms, resulting in a dose of 49.45 electrons per Å^2^. For saposin_ELIC_, spMSP1D1_ELIC_ and apo spMSP1D1_ELIC_, each movie consisted of 49 individual frames with a total dose of 47.55, 46.93 and 51.54 electrons per Å^2^, respectively.

### Single particle analysis and model building

Movies were motion corrected with MotionCor2^39^, and contrast transfer function (CTF) determined with GCTF v1.06^40^ using RELION3.1^41^. Particles were initially picked using LoG-based autopicking, followed by 2D class averaging to generate 2D classes for template-based picking. Particles were extracted and subjected to multiple rounds of 2D and 3D classification using a mask diameter of 140 Å. The initial model for 3D classification was generated using a 40 Å low-pass filtered map of MSP1E3D1_ELIC_ (EMD-27218), and 3D classifications and subsequent 3D refinements were performed using C5 symmetry. The best 3D refine map was then subject to post-processing, CTF refinement and Bayesian polishing.

An initial model of ELIC was obtained with MSP1E3D1_ELIC_ (pdb 8D66), which was used to perform real space refinement in PHENIX 1.19.2^42^. The structure was then manually built into the cryo-EM density map using COOT 0.9.6^43^ followed by iterative real space refinement in PHENIX and manual adjustments in COOT. Propylamine was fit in the density in the agonist binding site based on the predicted orientation of the amine group in cysteamine (another agonist of ELIC) from MD simulations^21^. To estimate the diameter of the nanodisc, unsharpened maps were low-pass filtered (8 Å) using relion image handler, as done by Noviello et *al*.^26^. The pointer atoms were placed at the edges and center of the map set at a contour of 1 σ and distances were measured using the distance tool in COOT.

### Modeling ELIC in a Nanodisc

To further examine the effect of nanodisc diameter on the conformation and dynamics of ELIC, we used molecular dynamics (MD) to simulate spMSP1D1_ELIC_ in two nanodiscs of differing size (9 nm and 11 nm). POPC was chosen as the sole lipid for these systems to avoid the uncertainty of placing different lipids in the model nanodisc, since significant lipid diffusion is not expected during the time scale of the simulation. By using the same starting structure of ELIC in both nanodisc systems, we can examine the effect of nanodisc size on ELIC structure. Starting with spMSP1D1_ELIC_, we modeled the missing residues (R318, G319, I320, T321, L322) assuming M4 remains an α-helix through its C-terminus. ELIC agonist, propylamine, was modeled into its binding site as in the cryo-EM structure. Parameters for propylamine were generated using the CGenFF server^44,45^. The local pKa of ionizable groups in ELIC side chains was determined with PROPKA3; this resulted in no protonation/deprotonation of any side chains in the protein with the system pH at 7.0. The N-terminus P11 was acetylated and the C-terminus was left charged as this is the true terminus. This structure was imported into CHARMM-GUI^46^ and placed in either an MSP1D1-33 (9 nm) nanodisc, MSP1E2D1 (11 nm) nanodisc, or planar bilayer^6^. The PPM server^47^ was used to orient ELIC such that the ion conduction pathway aligned to the normal of the plane delineated by the MSP bundle or membrane normal in the case of the planar bilayer system. The nanodisc and bilayer were composed of only POPC lipids to eliminate local lipid composition as a factor in conformational dynamics of ELIC. The ELIC-nanodisc construct was then solvated with the TIP3 water model ^48^ to provide ~2 nm of buffer between protein and the simulation box edge. Each system was then ionized with 150 mM NaCl and neutralized. The final simulation box measured 15.5×15.5×16.4 nm^3^ with 363,774 atoms for the 9 nm nanodisc simulation system, 17.0×17.0×17.1 nm^3^ with 457,197 atoms for the 11 nm nanodisc simulation system, and 12.1×12.0×14.5 nm^3^ with 162,265 atoms for the planar bilayer system.

Each simulation system was equilibrated for 50 ns. In each case, the backbone of all protein segments and the heavy atoms of phospholipids were harmonically restrained to their initial coordinates (k = 500 kcal/mol/nm^2^) for 5 ns. The harmonic position restraints were slowly removed over 20 ns with reduction of the harmonic force constant by 50 kcal/mol/nm^2^ every 2 ns. The system was then allowed 25 ns of equilibration without restraint, followed by 500 ns of unrestrained MD. Simulations were carried out in an NPT ensemble. Pressure was maintained at 1 atm using the Nosé-Hoover Langevin piston method^49,50^ with a piston period of 100 fs and piston decay of 50 fs. Temperature was maintained at 310 K with Langevin dynamics and a damping coefficient of 1 ps^-1^. A 2 fs timestep was used for integration. Short-range non-bonded interactions were cutoff after 12.0 Å with a switching function applied after 10.0 Å. Long-range electrostatics were handled using the particle mesh Ewald sums method^50^. MD simulation was carried out with NAMD3^51^. VMD^52^ was used for molecular visualization and quantitative analysis. The CHARMM36^53^ parameter set was used for lipids and ions with CHARMM36m^54^ and cation-π corrections^55^ being applied to protein segments.

### Analysis of ELIC in Nanodisc and Bilayer Systems

#### Nanodisc Diameter

It is known that nanodiscs do not retain a perfect circular shape, but demonstrate multiple conformational clusters that are ellipsoid^7,9^. Therefore, to demonstrate stability of nanodisc size in the 9 nm and 11 nm simulation systems, we generated an ellipse of best fit to the backbone atoms in both MSPs, as has been done previously^7^. The ellipse of best fit was recorded every 200 ps. The mean and standard deviation of nanodisc diameter was calculated over the last 250 ns of the simulation.

#### Membrane Thickness Measurement

To assess the effect of nanodisc size on the local membrane environment, we measured membrane thickness as a function of distance from the protein as well as two-dimensional thickness as a function of distance from the ion conduction pore; similar measurements have been made previously in empty nanodiscs^6,7^. Each frame of the simulation trajectory was aligned using the TMD of ELIC (residues 201 to 322). The instantaneous membrane midplane was taken to be the z-component of the geometric center of all phosphorous atoms in the system with lipids marked as being “upper” or “lower” leaflet depending on their position relative to the membrane midplane. All lipids in the system were then placed in a bin corresponding to the smallest distance between the phosphorous atom of each lipid and any TMD backbone atom. Bins were 0.3 nm wide and spanned from 0 nm to the radial width of the nanodisc (i.e. 4.5 nm for the 9 nm nanodisc system and 5.5 nm for the 11 nm nanodisc system). The instantaneous height of the bin was taken as the average height of all lipids within a bin, calculated using either the phosphorous atom or the geometric center of the glycerol backbone. The membrane thickness was then calculated as the distance between the height of upper lipids and lower lipids in each bin. The membrane thickness was measured every 200 ps over the last 250 ns of the simulation. For two-dimensional membrane thickness measurement, a similar process was used as above except binning was performed using polar coordinates (r,q) relative to the ion conduction pore with radial and theta bins at 5 Å and p/15 radian intervals, respectively. Again, membrane thickness was measured every 200 ps over the last 250 ns of the simulation.

#### Pore Radius Measurement

The functional states of pLGICs (*i*.*e*., resting, pre-active, activated, desensitized) are believed to have characteristic pore radius profiles that correspond to their ability to conduct ions^11^. In order to characterize how nanodisc diameter affects the structure of the ion conduction pore, we measured the pore radius for each simulation system as a function of time with the HOLE2^56^ module^57^ within MDAnalysis^58,59^. Each simulation system was aligned with its initial configuration, such that the ion conduction pore was oriented along the *z*-axis. Using a step size of 0.5 Å with a radius cut-off of 10 Å and the default van der Waal radii for atoms, the pore radius was measured every 200 ps over the course of the simulation trajectory.

## Data Availability

The data supporting the findings of this study are available within the paper and supplementary information files. The cryo-EM maps have been deposited in the Electron Microscopy Data Bank (EMDB) under accession codes EMD-28829 (SMA_ELIC_), EMD-28830 (saposin_ELIC_), EMD-28831 (spMSP1D1_ELIC_), and EMD-28832 (apo-spMSP1D1_ELIC_). The structural coordinates have been deposited in the RCSB Protein Data Bank (PDB) under the accession codes 8F32 (SMA_ELIC_), 8F33 (saposin_ELIC_), 8F34 (spMSP1D1_ELIC_), and 8F35 (apo-spMSP1D1_ELIC_).

## Acknowledgements

This study was supported by grants R35GM137957 to WC, F32GM139351 to JP, NSF2152059 to GB, and the Foundation for Anesthesia Education and Research Mentored Research Training Grant to MJA. We are grateful to Dr. Joe Henry Steinbach for helpful discussions.

## Authors Contributions Statement

W.W.C., J.T.P., V.D., M.J.A., and G.B. conceived the project and designed the experimental procedures. N.M.D. carried out the protein expression and purification. V.D. and J.T.P. prepared the nanodisc samples for cryo-EM and performed the single particle analysis. M.J.A. performed the MD simulations and analysis of the MD trajectories. M.R. and J.A.J.F. performed grid preparation and image acquisition for cryo-EM. All authors reviewed the paper.

## References

1. Denisov IG, Grinkova YV, Lazarides AA, Sligar SG. Directed self-assembly of monodisperse phospholipid bilayer Nanodiscs with controlled size. J Am Chem Soc 126, 3477–3487 (2004).

2. Flayhan A, Mertens HDT, Ural-Blimke Y, Martinez Molledo M, Svergun DI, Low C. Saposin Lipid Nanoparticles: A Highly Versatile and Modular Tool for Membrane Protein Research. Structure 26, 345–355 e345 (2018).

3. Orwick MC, et al. Detergent-free formation and physicochemical characterization of nanosized lipid-polymer complexes: Lipodisq. Angew Chem Int Ed Engl 51, 4653–4657 (2012).

4. Nasr ML, et al. Covalently circularized nanodiscs for studying membrane proteins and viral entry. Nat Methods 14, 49–52 (2017).

5. Zhang S, Ren Q, Novick SJ, Strutzenberg TS, Griffin PR, Bao H. One-step construction of circularized nanodiscs using SpyCatcher-SpyTag. Nat Commun 12, 5451 (2021).

6. Qi Y, Lee J, Klauda JB, Im W. CHARMM-GUI Nanodisc Builder for modeling and simulation of various nanodisc systems. J Comput Chem 40, 893–899 (2019).

7. Schachter I, Allolio C, Khelashvili G, Harries D. Confinement in Nanodiscs Anisotropically Modifies Lipid Bilayer Elastic Properties. J Phys Chem B 124, 7166–7175 (2020).

8. Sweeney DT, Krueger S, Sen K, Hackett JC. Structures and Dynamics of Anionic Lipoprotein Nanodiscs. J Phys Chem B 126, 2850–2862 (2022).

9. Bengtsen T, et al. Structure and dynamics of a nanodisc by integrating NMR, SAXS and SANS experiments with molecular dynamics simulations. Elife 9, e56518 (2020).

10. Kern DM, et al. Cryo-EM structure of SARS-CoV-2 ORF3a in lipid nanodiscs. Nat Struct Mol Biol 28, 573–582 (2021).

11. Howard RJ. Elephants in the Dark: Insights and Incongruities in Pentameric Ligand-gated Ion Channel Models. J Mol Biol 433, 167128 (2021).

12. Cheng WWL, Arcario MJ, Petroff JT. Druggable Lipid Binding Sites in Pentameric Ligand-Gated Ion Channels and Transient Receptor Potential Channels. Frontiers in Physiology 12, (2022).

13. Thompson MJ, Baenziger JE. Structural basis for the modulation of pentameric ligand-gated ion channel function by lipids. Biochim Biophys Acta Biomembr, 183304 (2020).

14. Yu J, et al. Mechanism of gating and partial agonist action in the glycine receptor. Cell 184, 957–968 e921 (2021).

15. Kim JJ, et al. Shared structural mechanisms of general anaesthetics and benzodiazepines. Nature 585, 303–308 (2020).

16. Laverty D, et al. Cryo-EM structure of the human alpha1beta3gamma2 GABAA receptor in a lipid bilayer. Nature 565, 516–520 (2019).

17. Rahman MM, et al. Structural mechanism of muscle nicotinic receptor desensitization and block by curare. Nat Struct Mol Biol 29, 386–394 (2022).

18. Zarkadas E, et al. Conformational transitions and ligand-binding to a muscle-type nicotinic acetylcholine receptor. Neuron 110, 1358–1370. e1355 (2022).

19. Petroff JT, et al. Open-channel structure of a pentameric ligand-gated ion channel reveals a mechanism of leaflet-specific phospholipid modulation. Nature Communications 13, 1–16 (2022).

20. Marabelli A, Lape R, Sivilotti L. Mechanism of activation of the prokaryotic channel ELIC by propylamine: a single-channel study. J Gen Physiol 145, 23–45 (2015).

21. Kumar P, et al. Cryo-EM structures of a lipid-sensitive pentameric ligand-gated ion channel embedded in a phosphatidylcholine-only bilayer. Proc Natl Acad Sci U S A 117, 1788–1798 (2020).

22. Kumar A, et al. Mechanisms of activation and desensitization of full-length glycine receptor in lipid nanodiscs. Nat Commun 11, 3752 (2020).

23. Zhuang Y, Noviello CM, Hibbs RE, Howard RJ, Lindahl E. Differential interactions of resting, activated, and desensitized states of the α7 nicotinic acetylcholine receptor with lipidic modulators. Proceedings of the National Academy of Sciences 119, e2208081119 (2022).

24. Dietzen NM, et al. Polyunsaturated fatty acids inhibit a pentameric ligand-gated ion channel through one of two binding sites. Elife 11, (2022).

25. Kinde MN, et al. Conformational Changes Underlying Desensitization of the Pentameric Ligand-Gated Ion Channel ELIC. Structure 23, 995–1004 (2015).

26. Noviello CM, et al. Structure and gating mechanism of the alpha7 nicotinic acetylcholine receptor. Cell 184, 2121–2134 e2113 (2021).

27. Zhang Y, et al. Asymmetric opening of the homopentameric 5-HT3A serotonin receptor in lipid bilayers. Nat Commun 12, 1074 (2021).

28. Basak S, Gicheru Y, Rao S, Sansom MS, Chakrapani S. Cryo-EM reveals two distinct serotonin-bound conformations of full-length 5-HT3A receptor. Nature 563, 270–274 (2018).

29. Du J, Lü W, Wu S, Cheng Y, Gouaux E. Glycine receptor mechanism elucidated by electron cryo-microscopy. Nature 526, 224–229 (2015).

30. Kasaragod VB, et al. Mechanisms of inhibition and activation of extrasynaptic αβ GABAA receptors. Nature 602, 529–533 (2022).

31. Polovinkin L, et al. Conformational transitions of the serotonin 5-HT3 receptor. Nature 563, 275–279 (2018).

32. Sauguet L, et al. Crystal structures of a pentameric ligand-gated ion channel provide a mechanism for activation. Proceedings of the National Academy of Sciences 111, 966–971 (2014).

33. Kumar P, Cymes GD, Grosman C. Structure and function at the lipid-protein interface of a pentameric ligand-gated ion channel. Proc Natl Acad Sci U S A 118, (2021).

34. Dacosta CJ, Dey L, Therien J, Baenziger JE. A distinct mechanism for activating uncoupled nicotinic acetylcholine receptors. Nature chemical biology 9, 701–707 (2013).

35. Hénault CM, et al. A lipid site shapes the agonist response of a pentameric ligand-gated ion channel. Nature chemical biology 15, 1156–1164 (2019).

36. Masiulis S, et al. GABAA receptor signalling mechanisms revealed by structural pharmacology. Nature 565, 454–459 (2019).

37. Nadezhdin KD, et al. Structural mechanism of heat-induced opening of a temperature-sensitive TRP channel. Nat Struct Mol Biol 28, 564–572 (2021).

38. Tong A, et al. Direct binding of phosphatidylglycerol at specific sites modulates desensitization of a ligand-gated ion channel. Elife 8, (2019).

39. Zheng SQ, Palovcak E, Armache JP, Verba KA, Cheng Y, Agard DA. MotionCor2: anisotropic correction of beam-induced motion for improved cryo-electron microscopy. Nat Methods 14, 331–332 (2017).

40. Zhang K. Gctf: Real-time CTF determination and correction. J Struct Biol 193, 1–12 (2016).

41. Zivanov J, et al. New tools for automated high-resolution cryo-EM structure determination in RELION-3. Elife 7, (2018).

42. Adams PD, et al. PHENIX: a comprehensive Python-based system for macromolecular structure solution. Acta Crystallogr D Biol Crystallogr 66, 213–221 (2010).

43. Emsley P, Lohkamp B, Scott WG, Cowtan K. Features and development of Coot. Acta Crystallogr D Biol Crystallogr 66, 486–501 (2010).

44. Vanommeslaeghe K, MacKerell Jr AD. Automation of the CHARMM General Force Field (CGenFF) I: bond perception and atom typing. Journal of chemical information and modeling 52, 3144–3154 (2012).

45. Vanommeslaeghe K, Raman EP, MacKerell Jr AD. Automation of the CHARMM General Force Field (CGenFF) II: assignment of bonded parameters and partial atomic charges. Journal of chemical information and modeling 52, 3155–3168 (2012).

46. Jo S, Kim T, Iyer VG, Im W. CHARMM-GUI: a web-based graphical user interface for CHARMM. Journal of computational chemistry 29, 1859–1865 (2008).

47. Lomize AL, Todd SC, Pogozheva ID. Spatial arrangement of proteins in planar and curved membranes by PPM 3.0. Protein Science 31, 209–220 (2022).

48. Jorgensen WL, Chandrasekhar J, Madura JD, Impey RW, Klein ML. Comparison of simple potential functions for simulating liquid water. The Journal of chemical physics 79, 926–935 (1983).

49. Martyna GJ, Tobias DJ, Klein ML. Constant pressure molecular dynamics algorithms. The Journal of chemical physics 101, 4177–4189 (1994).

50. Feller SE, Zhang Y, Pastor RW, Brooks BR. Constant pressure molecular dynamics simulation: The Langevin piston method. The Journal of chemical physics 103, 4613–4621 (1995).

51. Phillips JC, et al. Scalable molecular dynamics on CPU and GPU architectures with NAMD. The Journal of chemical physics 153, 044130 (2020).

52. Humphrey W, Dalke A, Schulten K. VMD: visual molecular dynamics. Journal of molecular graphics 14, 33–38 (1996).

53. Klauda JB, et al. Update of the CHARMM all-atom additive force field for lipids: validation on six lipid types. The journal of physical chemistry B 114, 7830–7843 (2010).

54. Huang J, et al. CHARMM36m: an improved force field for folded and intrinsically disordered proteins. Nature methods 14, 71–73 (2017).

55. Khan HM, MacKerell Jr AD, Reuter N. Cation-π interactions between methylated ammonium groups and tryptophan in the CHARMM36 additive force field. Journal of chemical theory and computation 15, 7–12 (2018).

56. Smart OS, Neduvelil JG, Wang X, Wallace B, Sansom MS. HOLE: a program for the analysis of the pore dimensions of ion channel structural models. Journal of molecular graphics 14, 354–360 (1996).

57. Stelzl LS, Fowler PW, Sansom MS, Beckstein O. Flexible gates generate occluded intermediates in the transport cycle of LacY. Journal of molecular biology 426, 735–751 (2014).

58. Michaud-Agrawal N, Denning EJ, Woolf TB, Beckstein O. MDAnalysis: a toolkit for the analysis of molecular dynamics simulations. Journal of computational chemistry 32, 2319–2327 (2011).

59. Gowers RJ, et al. MDAnalysis: a Python package for the rapid analysis of molecular dynamics simulations. In: Proceedings of the 15th python in science conference). SciPy Austin, TX (2016).

