## Supplementary Table and Figures for "Lipid nanodisc scaffold and size alters the structure of a pentameric ligand-gated ion channel"

**Supplementary Table 1: Cryo-EM data collection and refinement statistics**

| | $SMA_{ELIC}$ EMD-28829 PDB 8F32 | $saposin_{ELIC}$ EMD-28830 PDB 8F33 | $spMSP1D1_{ELIC}$ EMD-28831 PDB 8F34 | apo-$spMSP1D1_{ELIC}$ EMD-28832 PDB 8F35 |
|---|---|---|---|---|
| **Data collection and processing** | | | | |
| Magnification | 75000 | 96000 | 96000 | 96000 |
| Voltage (kV) | 300 | 300 | 300 | 300 |
| Electron exposure (e-/ Å$^2$) | 49.45 | 47.55 | 46.93 | 51.54 |
| Defocus range (μm) | -0.8 - 2.4 | -0.8 - 2.4 | -0.8 - 2.4 | -0.8 - 2.4 |
| Pixel size (Å) | 0.9 | 0.7 | 0.7 | 0.7 |
| Symmetry imposed | C5 | C5 | C5 | C5 |
| Initial particle images (no) | 1391288 | 1026975 | 1168080 | 837164 |
| Final particle images (no) | 128454 | 84696 | 65209 | 36562 |
| Map resolution (Å) | 3.96 | 3.50 | 3.32 | 3.38 |
| FSC threshold | 0.143 | 0.143 | 0.143 | 0.143 |
| **Refinement** | | | | |
| Initial model used | This study | This study | PDB 8D65 | This study |
| Model resolution (Å) | 3.80 | 3.47 | 3.40 | 3.30 |
| FSC threshold | 0.5 | 0.5 | 0.5 | 0.5 |
| Map sharpening B factor (Å$^2$) | -216.65 | -162.744 | -153.73 | -165.209 |
| **Model composition** | | | | |
| Non-hydrogen atoms | 12545 | 12545 | 12545 | 12525 |
| Protein Residues | 1535 | 1535 | 1535 | 1535 |
| Ligands | 5 | 5 | 5 | 0 |
| **B factors (Å$^2$) (0.5)** | | | | |
| Protein | 71.65 | 47.77 | 29.42 | 40.61 |
| Ligand | 20.00 | 20.00 | 20.00 | --------- |
| **R.m.s. deviations** | | | | |
| Bond lengths (Å) | 0.006 | 0.005 | 0.005 | 0.006 |
| Bond angles (°) | 1.145 | 1.097 | 1.113 | 1.148 |
| **Validation** | | | | |
| MolProbity Score | 1.85 | 1.82 | 1.50 | 1.60 |
| Clashscore | 5.80 | 5.88 | 2.12 | 2.73 |
| Poor rotamers (%) | 0.00 | 0.00 | 0.00 | 0.00 |
| **Ramachandran plot** | | | | |
| Favored (%) | 90.49 | 91.74 | 91.15 | 90.49 |
| Allowed (%) | 9.51 | 8.26 | 8.85 | 9.51 |
| Disallowed (%) | 0.00 | 0.00 | 0.00 | 0.00 |

### Supplementary Fig. 1

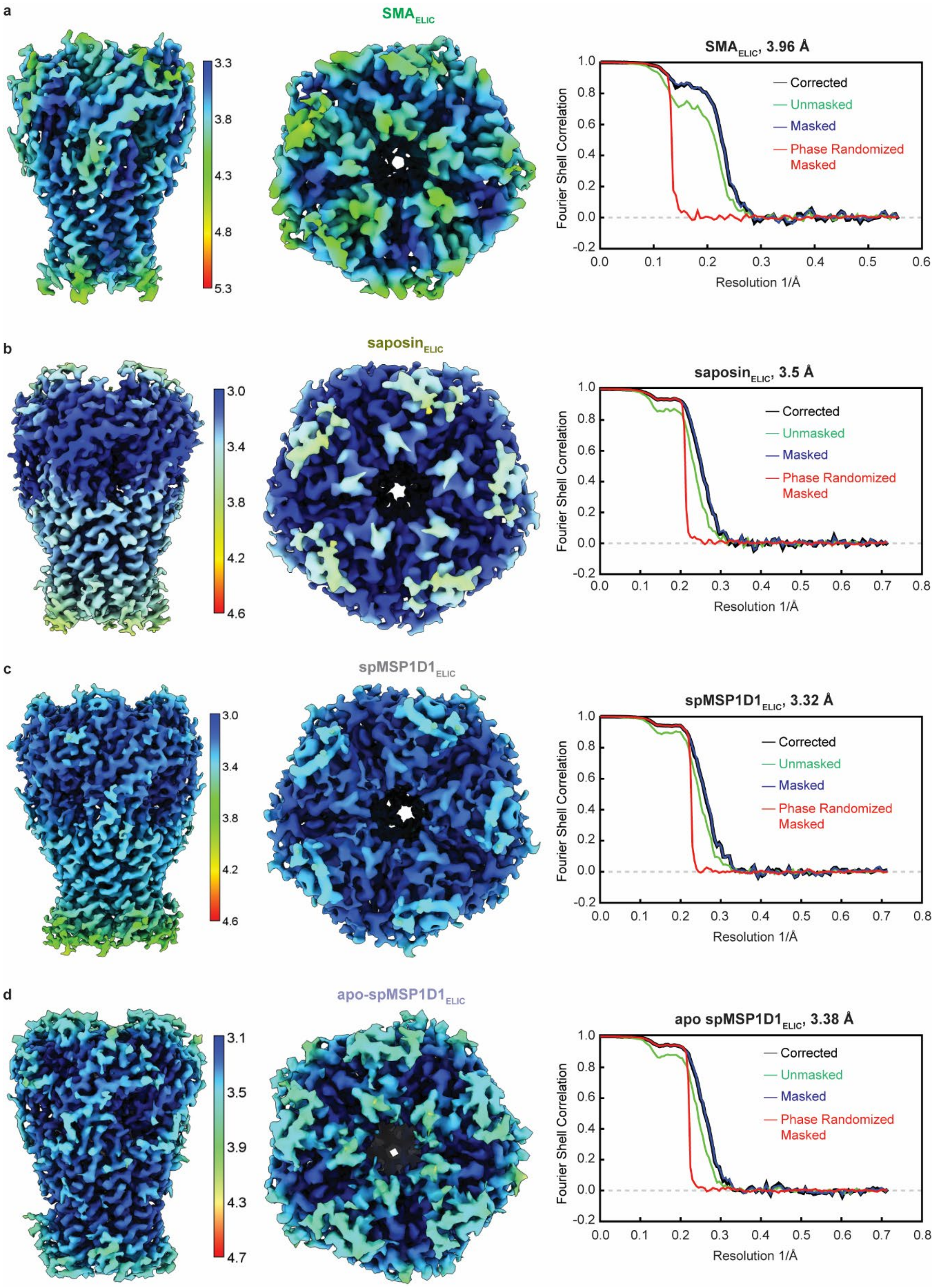

**<u>Supplementary Fig. 1:</u>** Post-processed maps colored according to local resolution and fourier shell correlation (FSC) curves for **(a)** $SMA_{ELIC}$, **(b)** $saposin_{ELIC}$, **(c)** $spMSP1D1_{ELIC}$, and **(d)** apo-$spMSP1D1_{ELIC}$. For each reconstruction, the local resolution was estimated by PHENIX using the sum of experimental half maps.

**Supplementary Fig. 2**

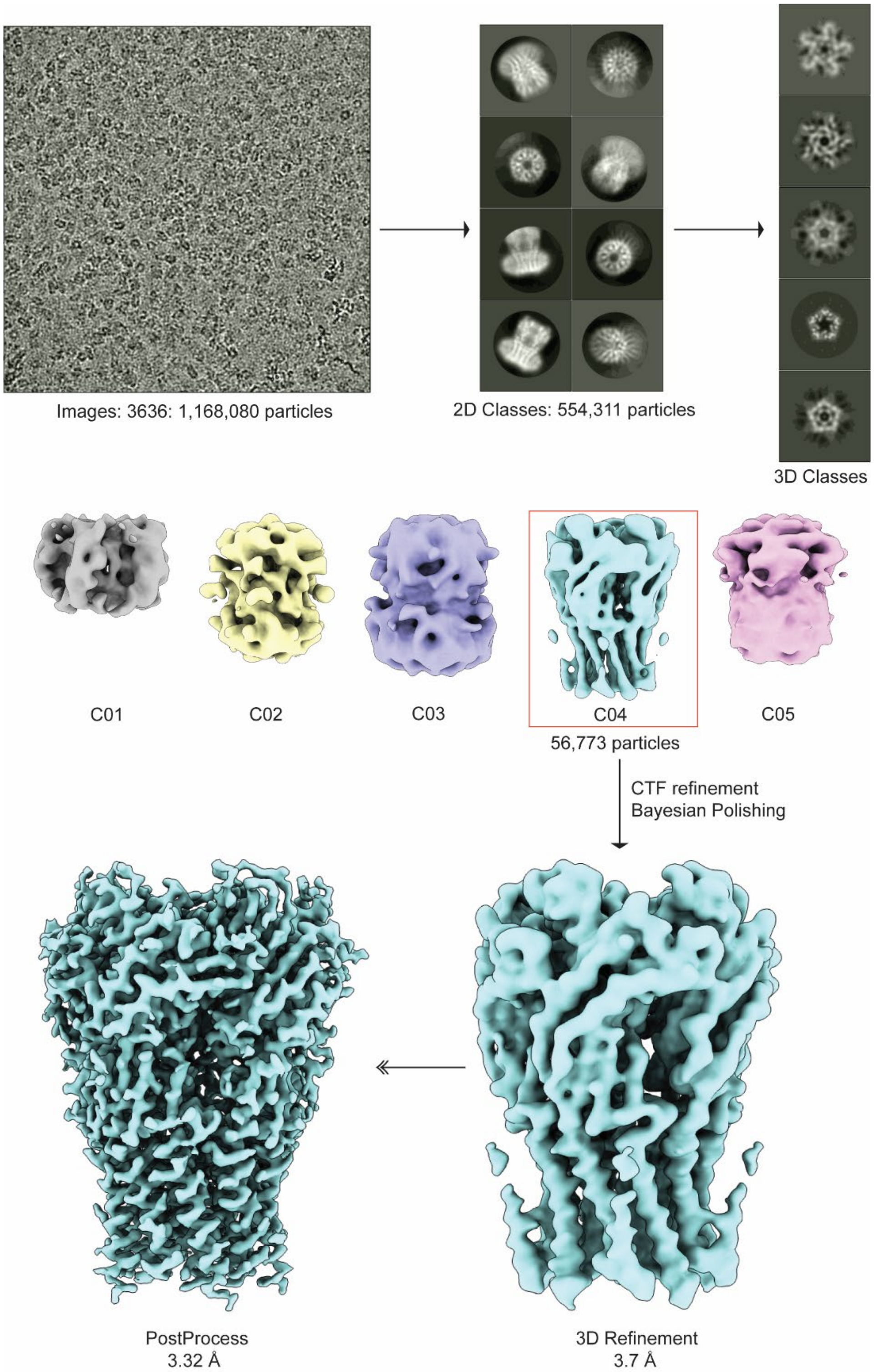


**Supplementary Fig. 2: Single particle cryo-EM 3D reconstruction of spMSP1D1$_{ELIC}$ using RELION3.1**. Workflow of single particle analysis for spMSP1D1$_{ELIC}$ showing a representative micrograph, 2D classes, 3D classes, and refined and post-processed maps. The same general approach was used to process the other structures ($SMA_{ELIC}$, $saposin_{ELIC}$, and apo-$spMSP1D1_{ELIC}$).

### Supplementary Fig. 3

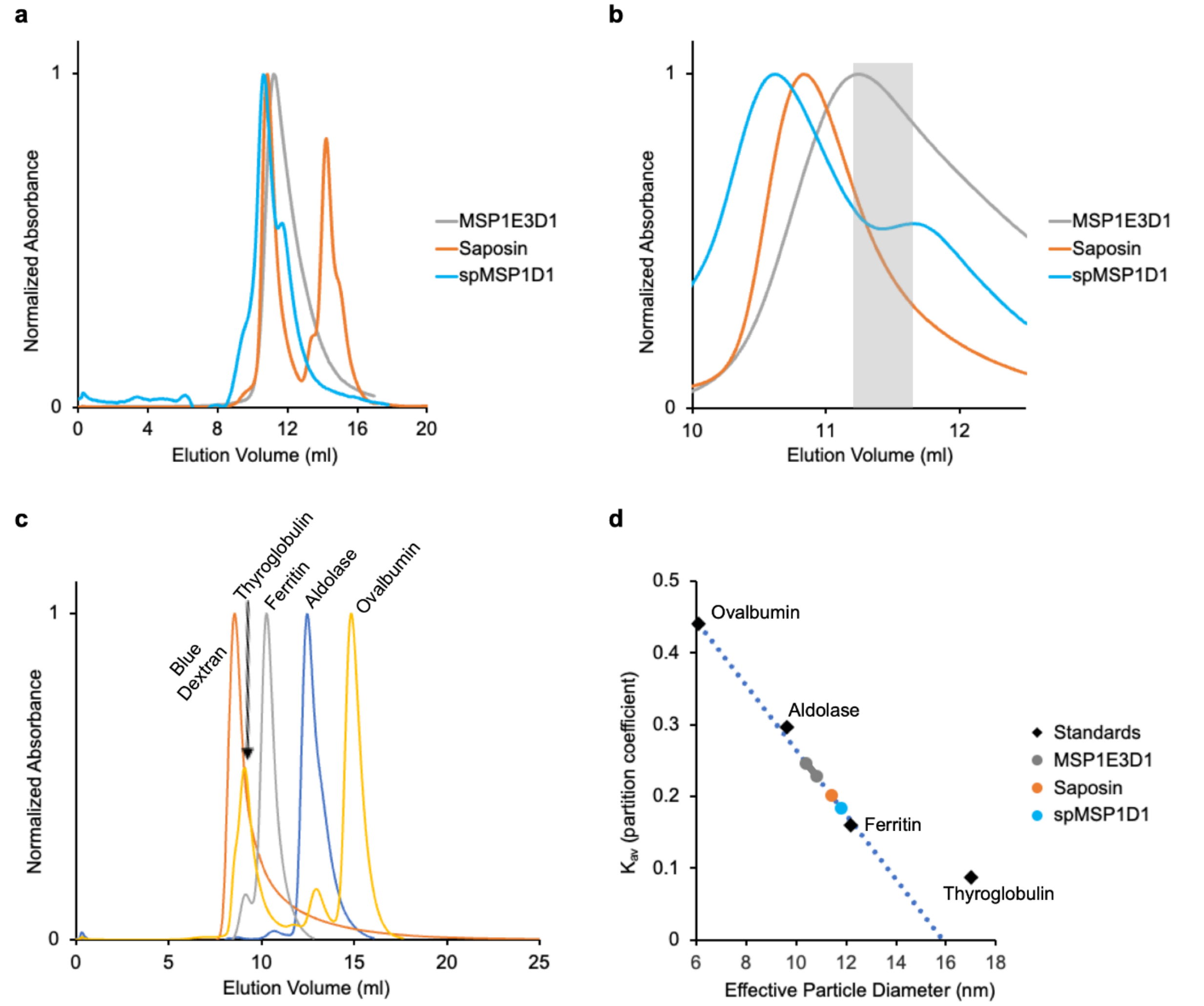


**e**

| | MSP1E3D1 | Saposin | spMSP1D1 |
|---|---|---|---|
| Kav | 0.247-0.266 | 0.201 | 0.184 |
| Eff Diameter (nm) | 10.4-10.8 | 11.4 | 11.8 |

**<u>Supplementary Fig. 3:</u> Size exclusion (SEC) analysis of ELIC in different nanodiscs (MSP1E3D1, saposin, and spMSP1D1) on Superdex 200 Increase 10/300 GL**. **(a)** and **(b)** SEC profiles of ELIC reconstituted in MSP1E3D1, saposin, and spMSP1D1. **(c)** SEC profile for standards: blue dextran, thyroglobulin, ferritin, aldolase, and ovalbumin. **(d)** Plot of partition coefficient (Kav) vs effective particle diameter (nm) for standards and ELIC in MSP1E3D1, saposin, and spMSP1D1. **(e)** Estimated effective particle diameter (nm) for ELIC in the indicated nanodiscs. The peak for ELIC in MSP1E3D1 was broad, see (b). Therefore, a range of elution volumes was used to estimate an effect particle diameter based on the eluted fractions used for cryo-EM.

Supplementary Fig. 4

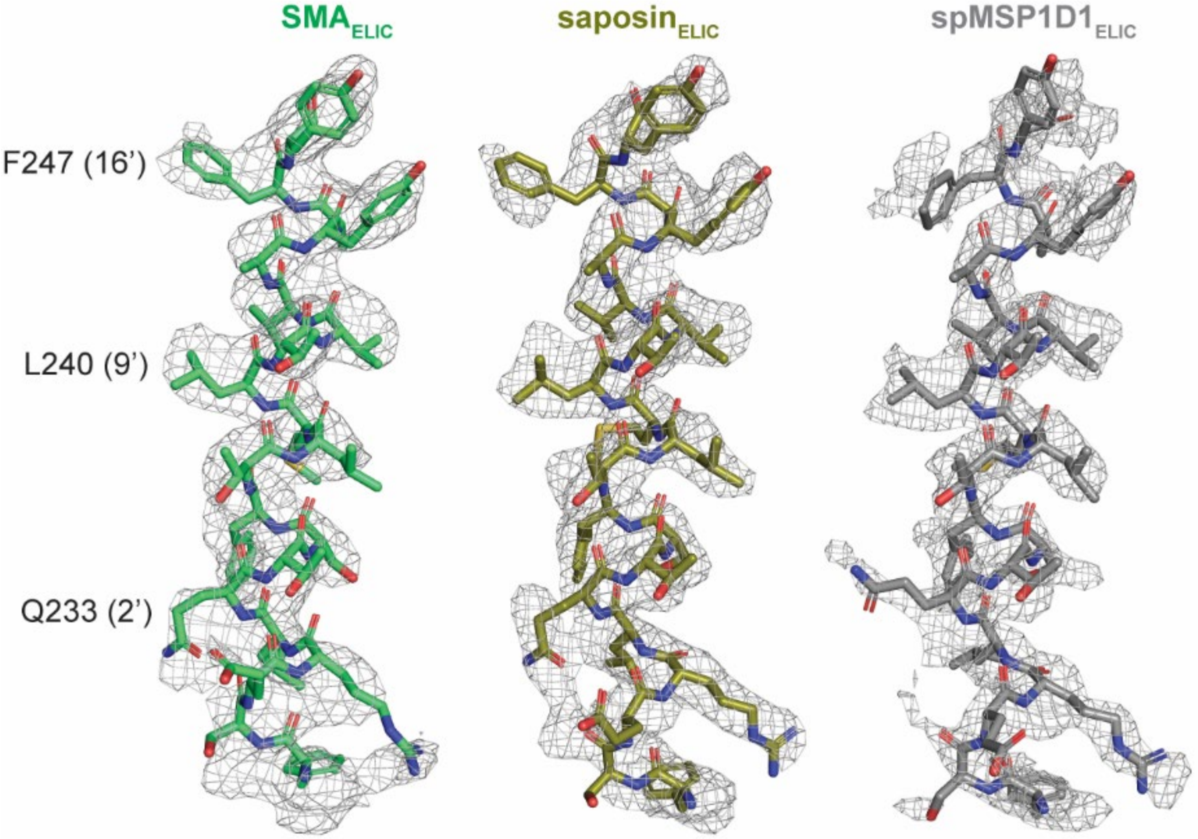


Supplementary Fig. 4: **Comparison of the cryo-EM density of M2 for $SMA_{ELIC}$, $saposin_{ELIC}$, and $spMSP1D1_{ELIC}$.** Density contours levels for $SMA_{ELIC}$, $saposin_{ELIC}$, and $spMSP1D1_{ELIC}$ are 1.8, 2.2, and 2.2 σ, respectively. The side chains for F247, L240 and Q233 are labeled. Molecular images were created in PyMOL.

**Supplementary Fig. 5**

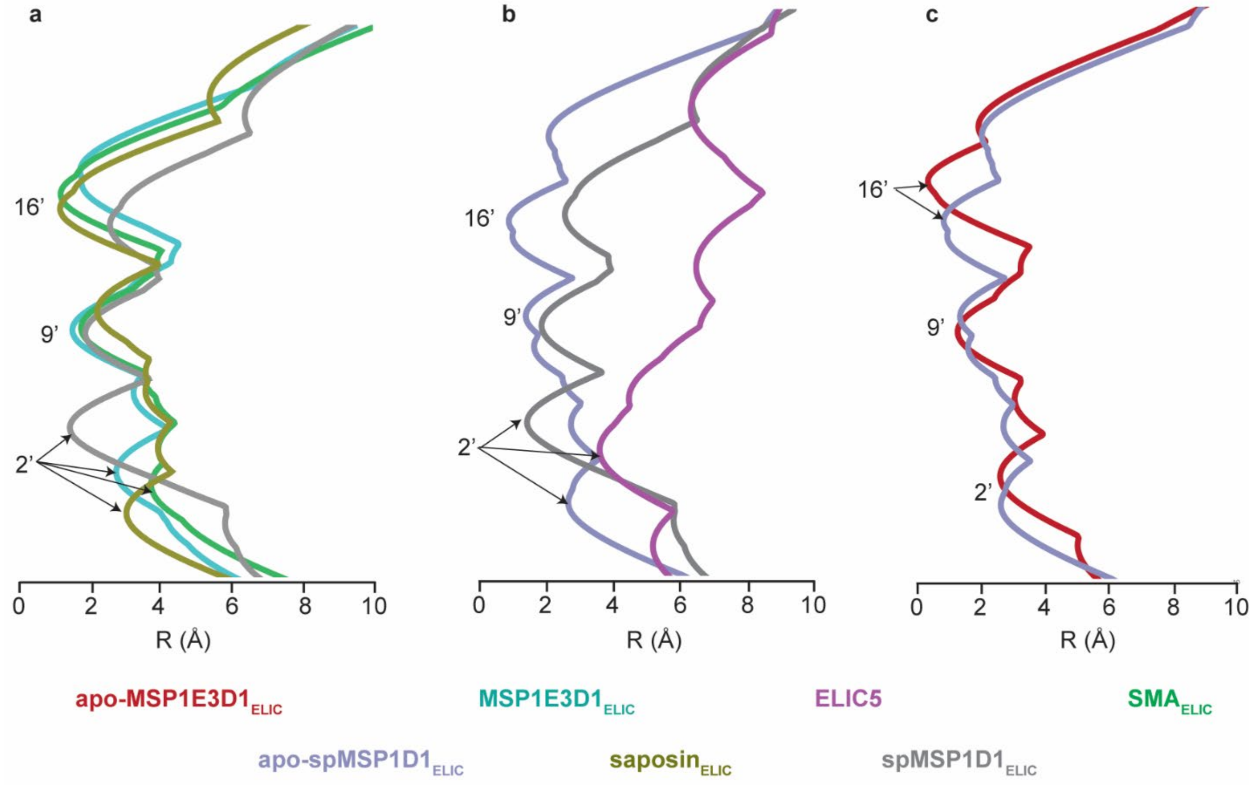


**<u>Supplementary Fig. 5:</u> Pore radius profiles for ELIC structures estimated by HOLE.** Pore radius profile comparison of **(a)** MSP1E3D1$_{ELIC}$, SMA$_{ELIC}$, saposin$_{ELIC}$, and spMSP1D1$_{ELIC}$, **(b)** spMSP1D1$_{ELIC}$, apo-spMSP1D1$_{ELIC}$, and ELIC5 (rotamer 2 for Q233), and **(c)** apo-MSP1E3D1$_{ELIC}$ and apo-spMSP1D1$_{ELIC}$. Overlapping profiles are represented from a global superposition of the structures.

**Supplementary Fig. 6**

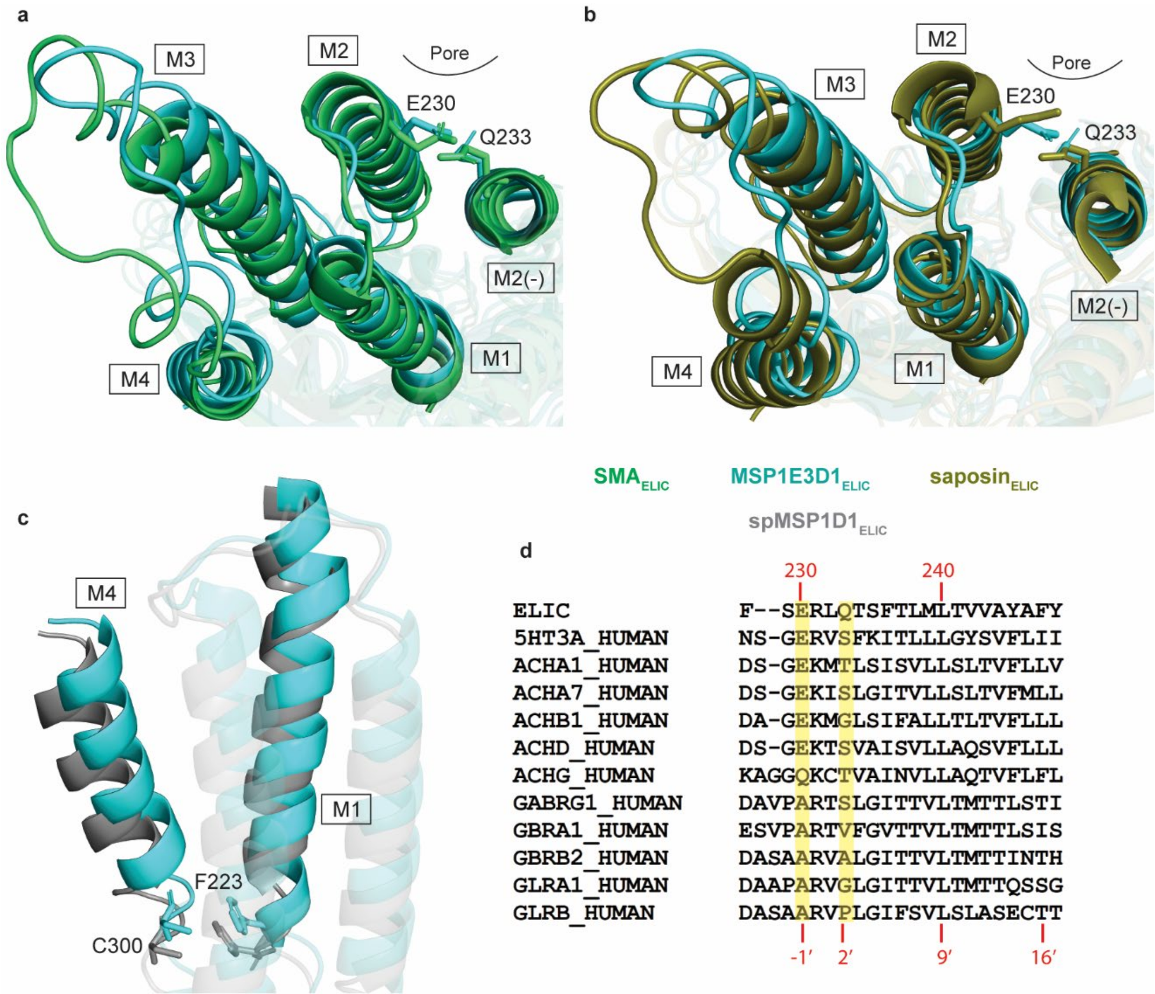


**Supplementary Fig. 6: TMD expansion in SMA$_{ELIC}$, saposin$_{ELIC}$, and spMSP1D1$_{ELIC}$.** Bottom-up view of the TMD of **(a)** SMA$_{ELIC}$ and MSP1E3D1$_{ELIC}$ and **(b)** saposin$_{ELIC}$ and MSP1E3D1$_{ELIC}$ showing the side chains of E230 and Q233. **(c)** Side view of M1 and M4 comparing spMSP1D1$_{ELIC}$ and MSP1E3D1$_{ELIC}$ and showing the side chains of C300 and F223. Images are from a global superposition of the indicated structures. **(d)** Sequence alignment of M2 in ELIC and other human pLGICs. E230 and Q233 of ELIC are highlighted in yellow. Other pLGIC sequences are from Uniprot, 5HT3A: P46098; ACHA1: P02708-2; ACHA7: P36544; ACHB1: P11230; ACHD: Q07001; ACHG: P07510; GABRG1:Q8N1C3; GBRA1: P14867; GBRB2: P47870; GLRA1: P23415; and GLRB: P48167.

**Supplementary Fig. 7**

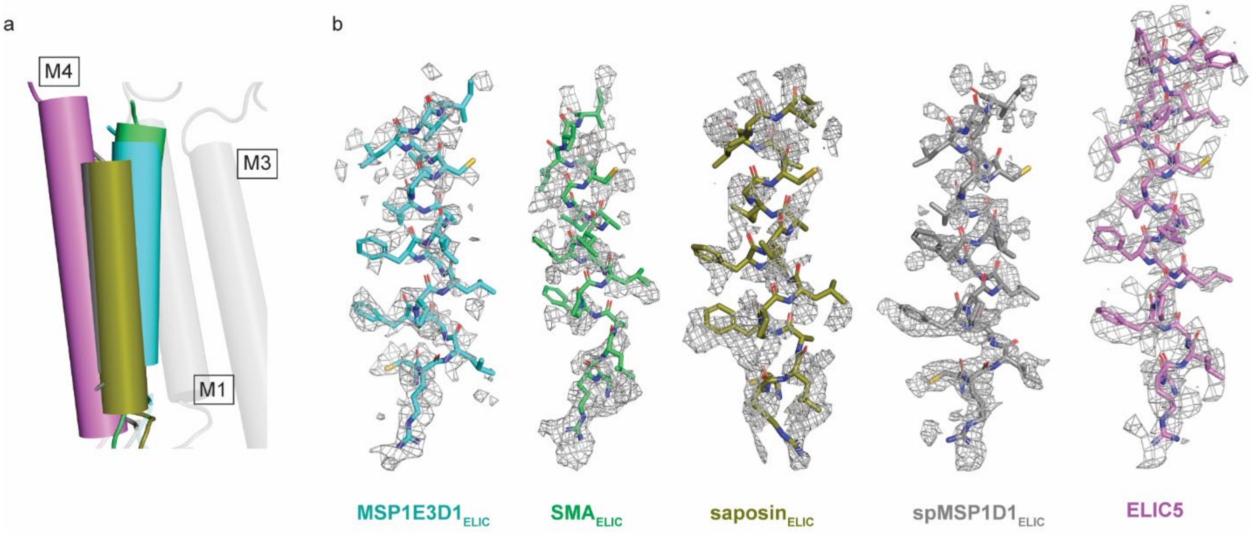


**Supplementary Figure 7: Structural (models and cryo-EM densities) comparison of M4 in MSP1E3D1$_{ELIC}$, spMSP1D1$_{ELIC}$, and ELIC5. (a)** Cylindrical model of M4 in the indicated structures, with a transparent M1 and M3 from spMSP1D1$_{ELIC}$ shown as a reference. Image shows a global superposition of the structures. In all structures except ELIC5, M4 was modeled up to I317 due to a lack of cryo-EM density in the C-terminus. The ELIC5 model extends to T321. **(b)** Cryo-EM density of M4 in the indicated structures contoured at 3.0 σ. Representations were generated in PyMOL.

**Supplementary Fig. 8**

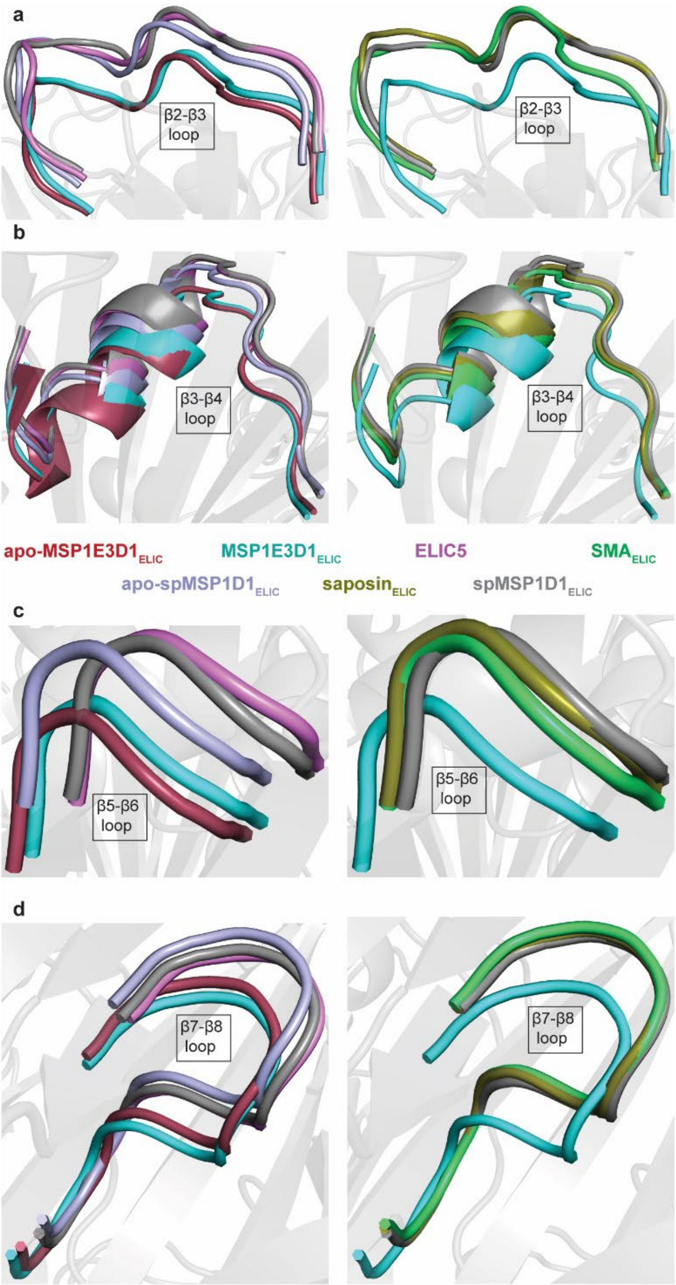


**Supplementary Fig. 8:** **Impact of nanodisc scaffold on the ECD.** Global superposition of indicated structures showing the **(a)** β2-β3, **(b)** β3-β4, **(c)** β5-β6, and **(d)** β7-β8 loops in the indicated structures. $MSP1E3D1_{ELIC}$ and apo-$MSP1E3D1_{ELIC}$ show a more compact or contracted ECD compared to ELIC5, $SMA_{ELIC}$, $saposin_{ELIC}$, $spMSP1D1_{ELIC}$, and apo-$spMSP1D1_{ELIC}$ that is evident by examining these ECD loops.

Supplementary Fig. 9

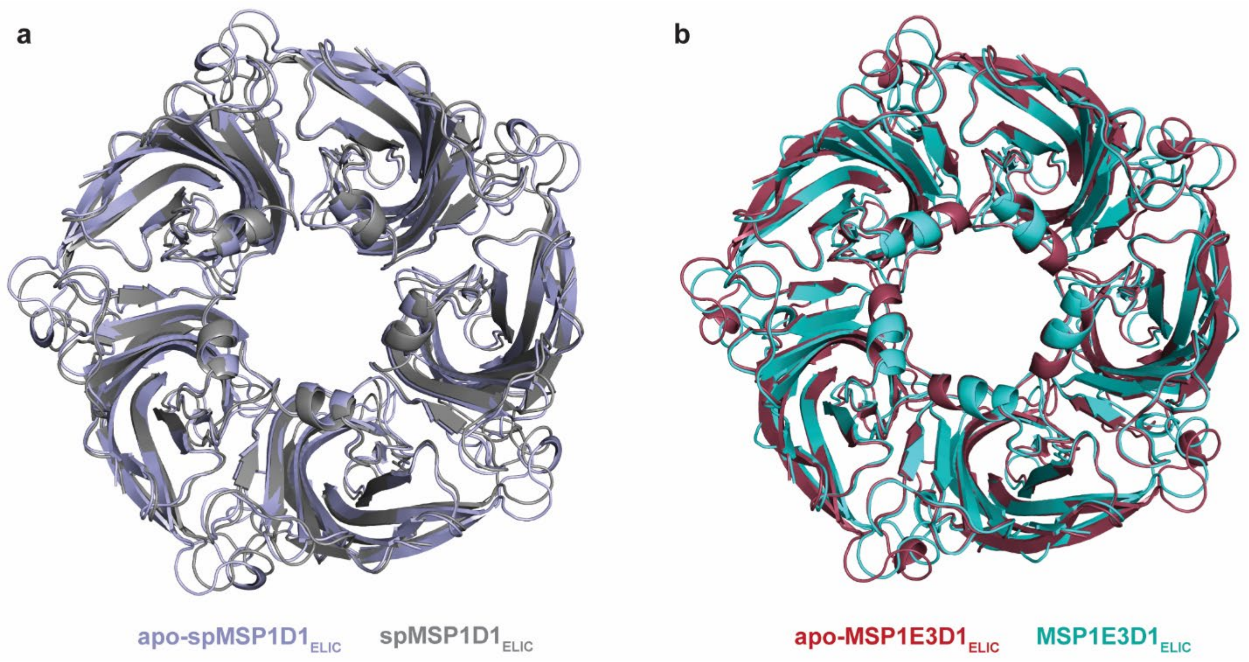


Supplementary Fig. 9: **ECD comparison of apo and agonist bound structures of ELIC in (a) spMSP1D1 or (b) MSP3E3D1 nanodiscs.** Representations show a global superposition of structures.

**Supplementary Fig. 10**

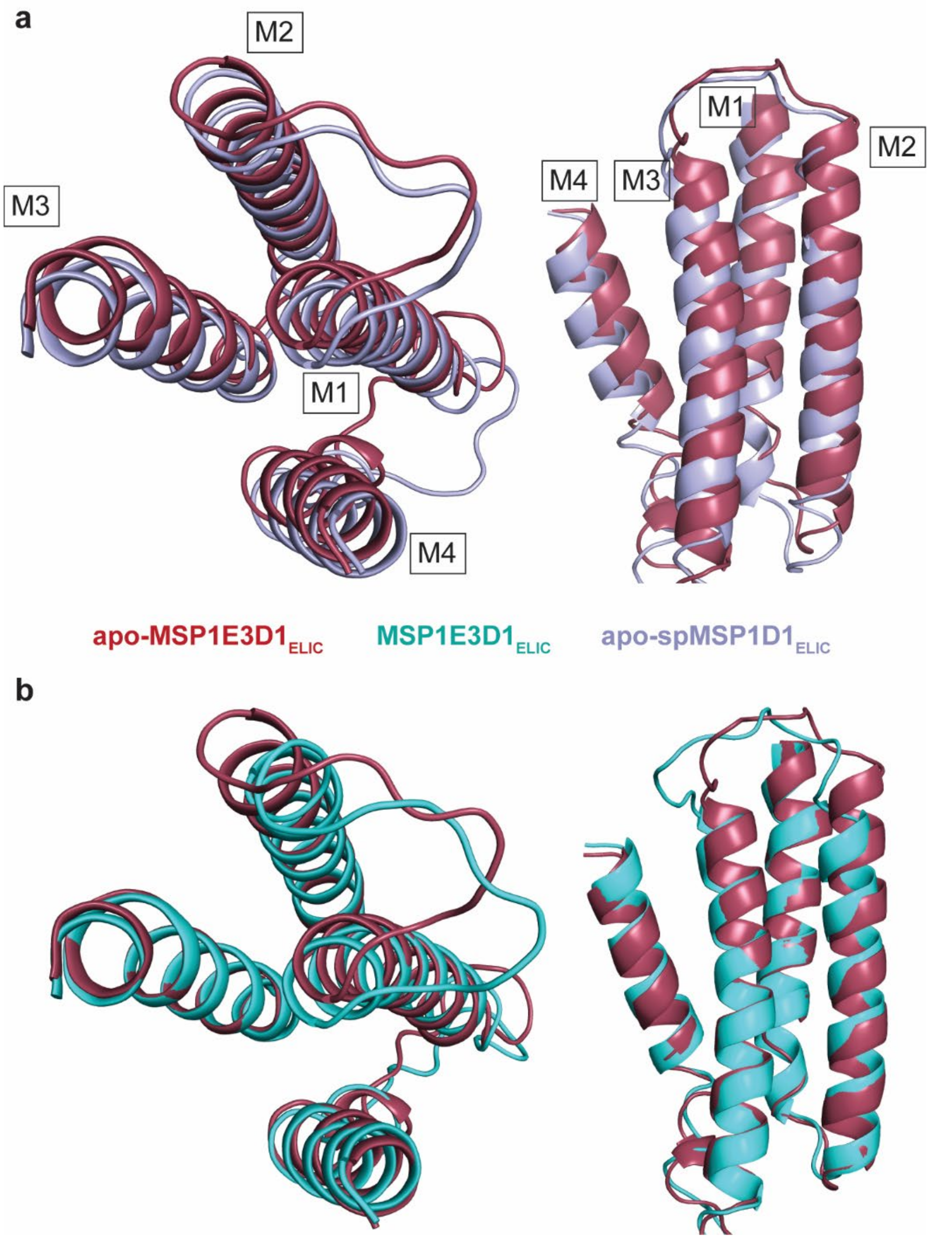


**Supplementary Fig. 10: Global superposition of TMD of apo-MSP1E3D1$_{ELIC}$, MSP1E3D1$_{ELIC}$, and apo-spMSP1D1$_{ELIC}$.** Top-down and side view of TMD comparing **(a)** apo-MSP1E3D1$_{ELIC}$ and apo-spMSP1D1$_{ELIC}$ and **(b)** apo-MSP1E3D1$_{ELIC}$ and MSP1E3D1$_{ELIC}$.

**<u>Supplementary Fig. 11</u>**

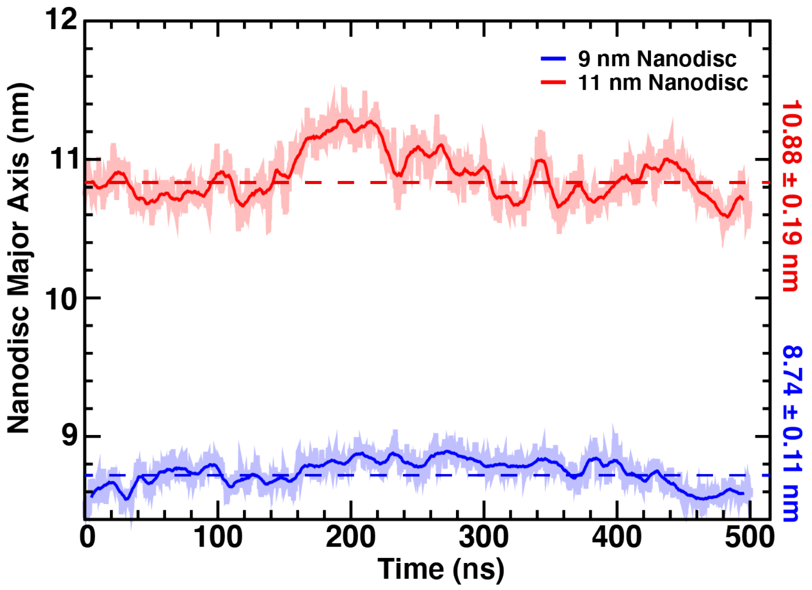


**<u>Supplementary Fig. 11:</u> Nanodisc major axis dimension as a function of simulation time.** The major axis of the 9 nm (blue) and 11 nm (red) nanodiscs are plotted throughout the course of the simulation with the location of the average size denoted by the dashed lines and labeled on the right of the plot (+/- 1 standard deviation). The major axis was measured at each time point by making an ellipse of best fit to all backbone atoms in each MSP. The heavy trace represents a running average over 25 measurements and the transparent trace represents the instantaneous measurement. The plot shows that the nanodisc diameter, as measured by the major axis, is stable at ~9 and 11 nm throughout the simulation.

**<u>Supplementary Fig. 12</u>**

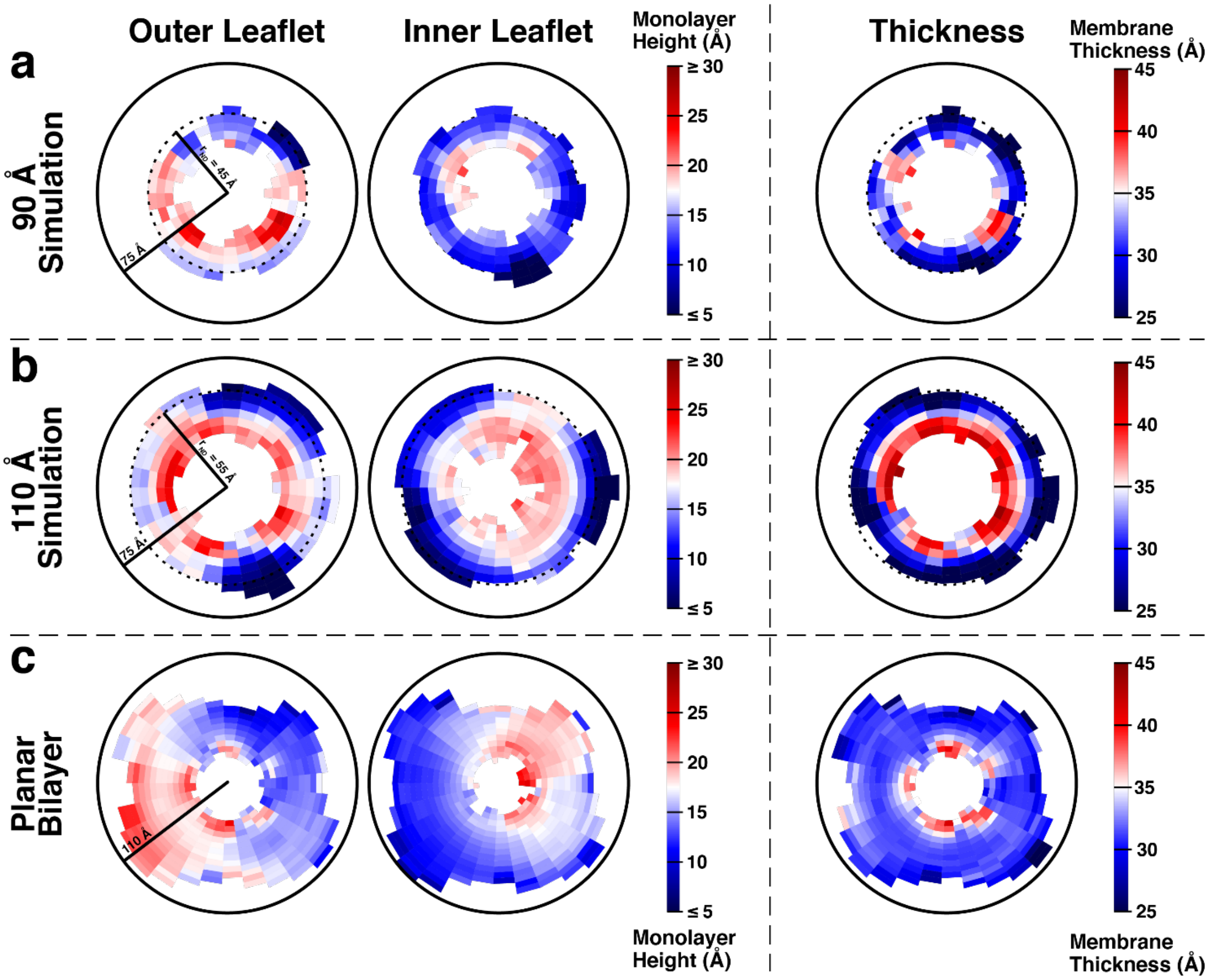


**<u>Supplementary Fig. 12:</u> Two-dimensional membrane thickness in nanodisc and bilayer.** The two-dimensional thickness of the membrane for the 9 nm nanodisc **(a)**, 11 nm nanodisc **(b)**, and lipid bilayer **(c)** simulations are displayed. In each simulation, the upper monolayer height (left), lower monolayer height (middle), and total membrane thickness (right) are shown. Membrane height is measured as phosphate-midplane distance in the case of monolayer height and phosphate-phosphate distance in the case of total membrane thickness. The monolayer plots have a different color bar scale than the total membrane thickness plots. Here, blue represents a thinner membrane, while red represents a thicker membrane. For the simulations containing a nanodisc, a circular dashed line displays, approximately, where the nanodisc rim would be located. This data was taken as an average over the last 250 ns of each simulation.
